# A global meta-analysis reveals higher variation in breeding phenology in urban birds than in their non-urban neighbours

**DOI:** 10.1101/2021.09.24.461498

**Authors:** Pablo Capilla-Lasheras, Megan J. Thompson, Alfredo Sánchez-Tójar, Yacob Haddou, Claire J. Branston, Denis Réale, Anne Charmantier, Davide M. Dominoni

**Author notes:** Corresponding author (PC-L). **Author contributions** PC-L, MJT, DR, AC and DMD conceived the study. PC-L, AS-T, CJB and DMD performed the literature search. PC-L extracted effect sizes from suitable published papers. MJT validated effect size extraction. PC-L and YH performed all statistical analysis with advice from AS-T. PC-L wrote the first draft of the manuscript with input from MJT, AS-T, DR, AC and DMD. All authors read and revised the manuscript. **Data availability** All R scripts and datasets needed to reproduce the analyses presented in this paper are available at: https://github.com/PabloCapilla/meta-analysis_variation_urban. Should the manuscript be accepted, a DOI to this data repository will be provided.

## Abstract

Cities pose a major ecological challenge for wildlife worldwide. Phenotypic variation, which can result from underlying genetic variation or plasticity, is an important metric to understand eco-evolutionary responses to environmental change. Recent work suggests that urban populations might have higher levels of phenotypic variation than non-urban counterparts. This prediction, however, has never been tested across species nor over a broad geographical range. Here, we conduct a meta-analysis of the avian literature to compare urban *versus* non-urban means and variation in phenology (i.e., lay date) and reproductive effort (i.e., clutch size, number of fledglings). First, we show that urban populations reproduce earlier and have smaller broods than non-urban conspecifics. Second, we show that urban populations have higher phenotypic variation in laying date than non-urban populations. This result arises from differences between populations within breeding seasons, conceivably due to higher landscape heterogeneity in urban habitats. These findings reveal a novel effect of urbanisation on animal life-histories with potential implications for species adaptation to urban environments (which will require further investigation). Higher variation in phenology in birds subjected to urban disturbance could result from plastic responses to a heterogeneous environment, or from higher genetic variation in phenology, possibly linked to higher evolutionary potential.

## Introduction

Humans have drastically changed environmental conditions on Earth, particularly since the invention of agriculture during the Neolithic Revolution. The footprint of human activity is most pronounced in urban environments, where microclimatic conditions, biogeochemical cycles and sensory landscapes are considerably different from those in non-urban habitats (Grimm *et al*. 2008). Perhaps not surprisingly, multiple shifts in animal and plant phenotypes have been associated with the novel conditions and selective pressures found in cities (Hendry *et al*. 2017). Indeed, numerous studies have reported divergent phenotypes between urban and non-urban populations in phenological, morphological, behavioural and reproductive traits (e.g., Alberti *et al*. 2017; Diamond *et al*. 2018; Campbell-Staton *et al*. 2020; reviewed in Johnson & Munshi-South 2017; Lambert *et al*. 2020; Diamond & Martin 2021). Most studies in urban ecology and evolution to date have focused on urban effects on *mean* phenotypes, and no study has explicitly investigated how urbanisation affects phenotypic *variation*. The extent to which populations can adapt to urban environments could be partly associated with how urbanisation affects their phenotypic variation (Thompson *et al*. 2022). Phenotypic variation is tightly linked to eco-evolutionary processes (Fusco 2001; Pavlicev *et al*. 2011): it is an essential condition for current selection, it results from past selection pressures, and it depends on gene flow and phenotypic plasticity. As such, assessing how urbanisation affects phenotypic variation can help us understand the potential for future phenotypic changes in urban environments and the eco-evolutionary implications of such changes (Thompson *et al*. 2022).

Recent single-species studies suggest that phenotypic variation could be affected by urbanisation (Caizergues *et al*. 2018; Gorton *et al*. 2018; Thompson *et al*. 2022). For example, in species with limited dispersal ability (i.e., whose dispersal occurs at a smaller scale than the scale at which the urban habitat varies), adaptation to local conditions could increase phenotypic variation within the urban matrix in heterogeneous urban environments. Findings from urban and non-urban meta-populations of the common ragweed (*Ambrosia artemisiifolia*) are consistent with this prediction as inter-population variation in several fitness proxies was greater in urban compared to non-urban environments (Gorton *et al*. 2018). A meta-analysis of selection strength found weaker selection occurring in human-disturbed populations (Fugère & Hendry 2018; note that this analysis did not specifically test the effect of urbanisation on selection strength and only included one study directly associated with urbanisation), which if extrapolated to the urban context, could lead to higher phenotypic variation in urban populations compared to their non-urban counterparts. Overall, these studies converge with the notion that urban populations could display higher levels of phenotypic variation due to several eco-evolutionary processes. These findings also highlight that the extent to which urbanisation might impact phenotypic variation likely depends on the interplay between the temporal and spatial scale at which environmental conditions fluctuate in the urban habitat, as well as on the species’ longevity and dispersal ability (Thompson *et al*. 2022).

The temporal scale at which differences in phenotypic variation between urban and non-urban habitats manifest can help us evaluate their ecological causes, and is likely to determine the eco-evolutionary implications of increased phenotypic variation in urban habitats (Thompson *et al*. 2022). First, urban populations could display higher phenotypic variation than non-urban populations within a given breeding season (i.e., intra-annual variation; as a result, for example, of consistent differences in landscape heterogeneity between habitats; Pickett et al. 2017). Second, urban populations could display higher phenotypic variation than non-urban populations due to larger yearly fluctuations in environmental conditions (i.e., inter-annual variation; if, for example, urban populations are more sensitive to changes in weather), with or without intra-annual differences in phenotypic variation between urban and non-urban populations. In the latter scenario, similar levels of phenotypic variation would be exposed to natural selection in short-lived species (e.g., annual species).

Urban environments have been referred to as spatially more heterogeneous than non-urban habitats of the same geographical area (Pickett *et al*. 2017). High urban habitat heterogeneity could increase phenotypic variation compared to adjacent non-urban habitats if, for example, urban organisms change their phenotype according to local environmental conditions (e.g., through either developmental or later-life phenotypic plasticity). The empirical assessment of this idea, however, largely depends on the scale at which urban habitat heterogeneity is measured, the spatial scale at which the organism of interest operates and the heterogeneity of the non-urban habitat of reference (Pickett *et al*. 2017; Uchida *et al*. 2021). For example, a megacity could be spatially heterogeneous, containing a diverse array of habitats (e.g., multiple urban parks with different ecological conditions, a varying level of impervious surface, etc.), and, thus, be overall vastly more heterogeneous than a neighboring non-urban habitat. However, species could reduce the range of environmental conditions that they experience through matching habitat choice (e.g., Muñoz et al. 2014), limiting the potential effect of urban habitat heterogeneity on phenotypic variation. Therefore, measuring habitat heterogeneity at different spatial scales will be paramount to understand the potential association between habitat heterogeneity and increased phenotypic variation in urban areas.

Here, we investigate how urbanisation impacts mean phenotypic values and phenotypic variation using a meta-analysis of 399 paired urban and non-urban comparisons of avian life-history traits (laying date, clutch size and number of fledglings) published between 1958 and 2020 including 35 bird species (Figure 1). We use paired within species urban – non-urban comparisons to investigate the following questions: i) Is urbanisation associated with shifts in mean life-history traits? ii) Is urbanisation associated with changes in variation in life-history traits? iii) What is the temporal and spatial scale at which urbanisation correlates with changes in phenotypic variation? Based on previous research (Chamberlain *et al*. 2009; Sepp *et al*. 2018), we predict that urban bird populations display on average earlier phenology, smaller clutch size and lower number of fledglings than non-urban populations. We also predict increased phenotypic variation in urban populations compared to non-urban populations for all three traits examined (see above). We disentangle urban effects on phenotypic variation across different temporal and spatial scales, suggesting an ecological mechanism for the effects of urbanisation on avian phenotypic variation. This study provides, for the first time, meta-analytical evidence that urban conditions can magnify phenotypic variation in phenology and highlights the potential role of increased habitat heterogeneity in urban areas as an ecological mechanism underlying this effect.

**Figure 1.**
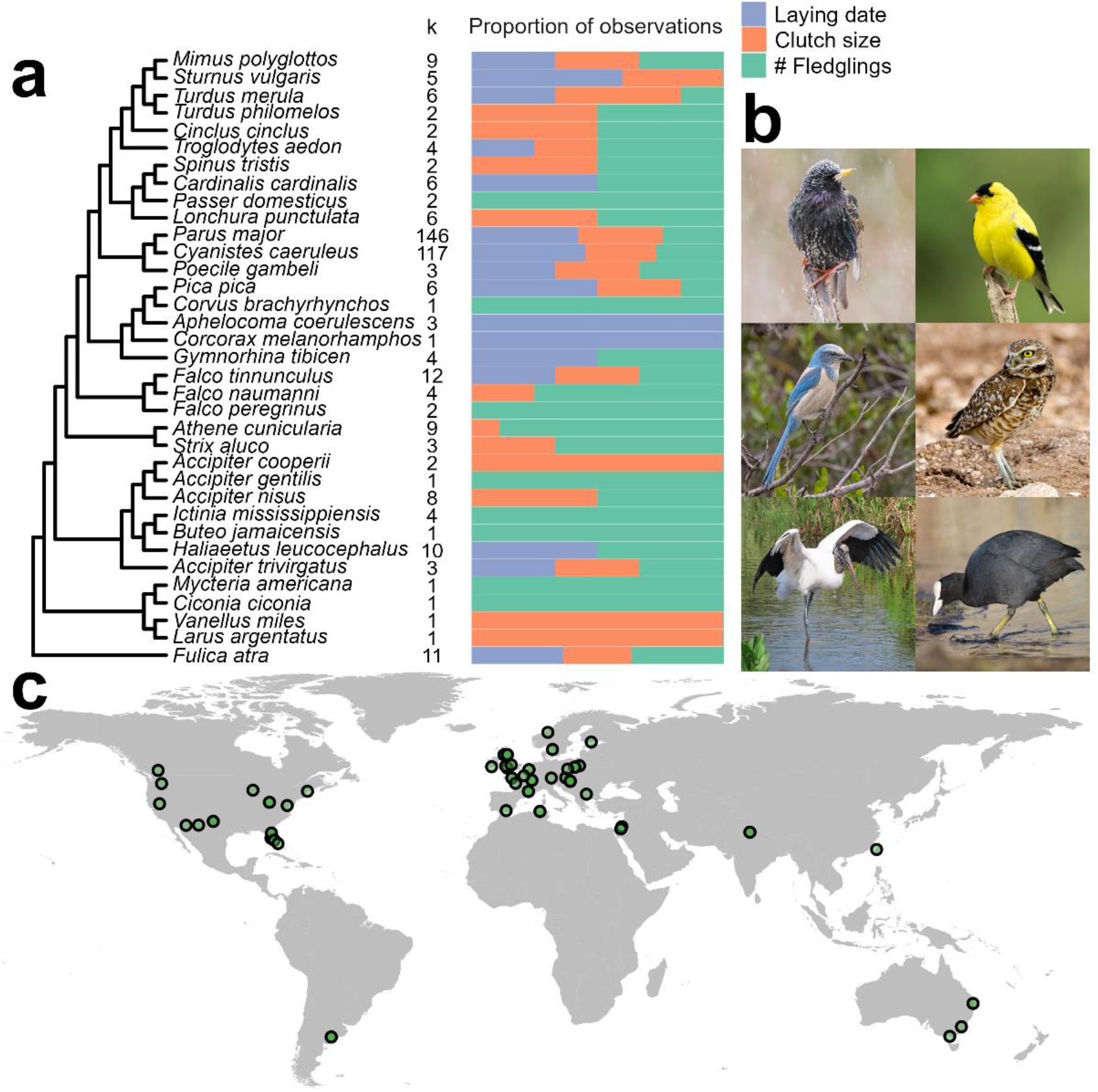
Phylogenetic and geographical breadth of the meta-analytic data. **(a)** Phylogenetic tree of the 35 avian species included in the meta-analysis along with the number of effect sizes (i.e., urban – non-urban comparisons) included per species (‘k’; which may encompass multiple years of study from the same publication) and the proportion of observations for each life-history trait (purple: Laying date; orange: Clutch size; Green: Number of fledglings). (**b**) Our meta-analysis included a broad range of species, as examples, left to right from top to bottom: *Sturnus vulgaris, Spinus tristis, Aphelocoma coerulescens, Athene cunicularia, Mycteria americana* and *Fulica atra*. All images are copyright free (CC - Public Domain Mark 1.0. Authors: Shenandoah National Park [first two images], Mike Carlo/U.S. Fish & Wildlife Service, Jennifer Soos, Susan Young and Ekaterina Chernetsova) and were extracted from www.flickr.com. (**c**) Global map (excluding Antarctica) showing the location of each study included in the meta-analysis. Each point represents one study area in which one or more urban – non-urban pairs of populations were sampled across a varying number of years.

## Material and methods

### Literature review

We began our literature search by inspecting two published reviews on the impact of urbanisation on avian biology (Chamberlain *et al*. 2009; Sepp *et al*. 2018). As we were interested in how phenology and reproduction were affected by urbanisation, we identified studies cited in Chamberlain *et al*. (2009) (n = 37) and Sepp *et al*. (2018) (n = 32) that could contain either raw data, or mean and variance estimates for first clutch laying initiation (hereafter laying date), clutch size and number of nestlings fledged per breeding attempt (hereafter number of fledglings), for paired urban and non-urban populations (see details below). Then, we performed four searches of the Web of Science Core Collection on the 27^th^ of October 2020 (databases covered: SCI-EXPANDED – 1900-present, SSCI – 1956-present, A&HCI – 1975-present, BKCI-S – 2005-present, BKCI-SSH – 2005-present and ESCI – 2015-present) to recover studies published since 1900 and including all languages and all document types. We performed the following four searches on the Web of Science Core Collection: (**1**) TS=(“urban*” AND (“bird*” OR “aves” OR “avian” OR “ornithol*” OR “passerine*” OR “passeriform*” OR “songbird*” OR *list of bird genera*) AND (“laying date” OR “lay date” OR “first egg” OR “clutch size” OR “eggs laid” OR “number of eggs” OR “fledgling*” OR “fledging” OR “reproductive success” OR “fitness”)); (**2**) TS=(“urban*” AND “bird” AND “laying date”); (**3**) TS=(“urban*” AND “bird” AND “clutch size”); (**4**) TS=(“urban*” AND “bird” AND “fledglings”). The *list of avian genera* in the first search string consisted of a list of all avian genera and can be found in Supplementary text D (see also acknowledgements). We complemented the search on the Web of Science Core Collection by searching Scopus using search string ‘(1)’ above (Scopus field ‘TITLE-ABS-KEY’). Both literature searches, on the Web of Science Core Collection and Scopus, included studies published before the 27^th^ of October 2020. We used the literature search results in these two major search engines to assess the comprehensiveness of our search (see Supplementary Text A for details). These searches found 892, 71, 198, 167 (on the Web of Science Core Collection) and 735 (on Scopus) studies, respectively, which we combined with the studies identified from Chamberlain *et al*. (2009) and Sepp *et al*. (2018) to create a list of 2,132 (non-unique) studies (Figure S1). We then de-duplicated this list using the R package ‘revtools’ (using exact matching of study titles in function ‘*find_duplicates*’, v0.4.1; Westgate 2019) and by manually inspecting all titles and author lists. Our final list contained 1,166 unique studies (Figure S1), which we screened by reading their title and abstract (this first screening step was made by PC-L, CJB and DMD). If the title and/or abstract indicated that the paper could fit our requirements for data collection (see below), we read the study fully, aiming to extract mean, standard deviation (SD) and sample size (n) of our life-history traits of interest for urban and non-urban bird populations. If SD was not available but authors provided SE, the former was calculated as: *SD* = *SE* × √*n*. Mean and SD were extracted from data quartiles and medians in four effect sizes from two studies following (Luo *et al*. 2016; Shi *et al*. 2020). When available, we extracted estimates per breeding season (i.e., papers sometimes reported mean, SD and n for urban and non-urban populations in multiple breeding seasons). If a study reported incomplete information for inclusion in our meta-analysis (e.g., mean was provided but not SD or SE), we contacted the authors to ask for this missing information (a complete list of authors that provided estimates can be found in the acknowledgements).

### Criteria for inclusion

We were interested in investigating the effects of urbanisation on life-history traits, with an interest in testing the association between urbanisation and, mean and variation in trait values. Paired urban – non-urban designs, where an urban population is compared to an adjacent non-urban population, are a powerful approach to identify the effects of urban living while controlling for temporal and geographical variation, and large-scale genetic structure among populations (Caizergues *et al*. 2021; Salmón *et al*. 2021). Therefore, we included studies if they compared geographically close (i.e., paired) urban and non-urban populations and reported laying date of the first clutches of the season, clutch size or number of fledglings for the same breeding season across both habitats. When multiple populations were compared along a gradient of urbanisation, we extracted estimates for the two populations at the extremes of the gradient (i.e., most and least urbanised populations). When studies combined estimates across several breeding seasons, we included them in our meta-analysis if urban and non-urban populations had been sampled in the same breeding seasons. All effect sizes were extracted by one author (PC-L). To validate data extraction, another author (MJT) checked 15% of the studies included in the meta-analysis, comprising 55 effect sizes (17.80% of the final data set; Supplementary Text B).

Initially, our dataset contained 443 paired urban – non-urban estimates from 40 bird species and 74 studies. Of these, three observations were removed due to missing sample sizes, 26 observations were removed due to missing SD and 11 observations were removed because their sample size was one (which precludes the calculation of mean and SD). Four observations were removed because they reported a SD of zero (these indeed had very low sample sizes: 3, 2, 7, 2 observations). Our final dataset included 399 comparisons between paired urban – non-urban populations from 35 bird species and 68 studies (Figure 1; refs.: Middleton 1979; Schmidt & Steinbach 1983; Dhondt *et al*. 1984; Eden 1985; Stout *et al*. 1998; Boal & Mannan 1999; Mcgowan 2001; Schoech & Bowman 2001; Solonen 2001, 2014; Antonov & Atanasova 2003; Rollinson & Jones 2003; Liven-Schulman *et al*. 2004; Millsap *et al*. 2004; Sharma *et al*. 2004; Beck & Heinsohn 2006; Conway *et al*. 2006; Mennechez & Clergeau 2006; Charter *et al*. 2007; Isaksson & Andersson 2007; Kelleher & O’Halloran 2007; Schoech *et al*. 2007; Hinsley *et al*. 2008; Isaksson *et al*. 2008; Newhouse *et al*. 2008; Solonen & Ursin 2008; Berardelli *et al*. 2010; Ibáñez-Álamo & Soler 2010; Shustack & Rodewald 2011; Seress *et al*. 2012, 2018, 2020; Stracey & Robinson 2012; Brahmia *et al*. 2013; Cardilini *et al*. 2013; Morrissey *et al*. 2014; Sumasgutner *et al*. 2014; Gahbauer *et al*. 2015; Glądalski *et al*. 2015, 2016b, a, 2017, 2018; Lin *et al*. 2015; Wawyrzyniak *et al*. 2015; Bailly *et al*. 2016; Minias 2016; Perlut *et al*. 2016; Biard *et al*. 2017; Capilla-Lasheras *et al*. 2017; Kopij 2017; Lee *et al*. 2017; Pollock *et al*. 2017; Preiszner *et al*. 2017; Thornton *et al*. 2017; Bobek *et al*. 2018; Caizergues *et al*. 2018; Gryz & Krauze-Gryz 2018; de Satgé *et al*. 2019; Hajdasz *et al*. 2019; Kettel *et al*. 2019; Rosenfield *et al*. 2019; Welch-Acosta *et al*. 2019; Baldan & Ouyang 2020; Evans & Gawlik 2020; Jarrett *et al*. 2020; Luna *et al*. 2020; Partecke *et al*. 2020). Of these 399 comparisons, 151 corresponded to comparisons of laying date (n = 32 studies), 119 were comparisons of clutch size (n = 42 studies) and 129 were comparisons of number of fledglings (n = 48 studies) (Figure S2). Last, there were 363 comparisons for single years (n = 47 studies) and an additional 36 comparisons included estimates across multiple years (n = 21 studies).

### Meta-analytic effect sizes

We standardised laying date across studies by coding it as the number of days after the 1^st^ of January (January 1^st^ = 1). Mean laying date estimates across habitats always fell within the same calendar year. For each of the three life-history traits, we computed the log response ratio (lnRR) and the log coefficient of variation ratio (lnCVR) to investigate differences in mean values and variability between urban and non-urban populations (Hedges *et al*. 1999; Nakagawa *et al*. 2015; Senior *et al*. 2020). We calculated lnRR and lnCVR along with their associated sampling variances (Nakagawa *et al*. 2015) using the R function ‘*escalc*’ in the ‘metafor’ R package (v3.4.0; Viechtbauer 2010). Both lnRR and lnCVR were calculated so that positive values meant higher estimates in urban populations compared to their non-urban counterparts. Often mean and variance values are positively associated (e.g., Taylor’s Law; (Nakagawa & Schielzeth 2013; Cohen & Xu 2015). Therefore, we chose lnCVR over lnVR (i.e., log total variation ratio; Nakagawa *et al*. 2015) as the former accounts for the mean– variance relationship (Nakagawa *et al*. 2015; Senior *et al*. 2020). However, we carried out sensitivity analysis using, among others, the log total variation ratio (section ‘Sensitivity analyses’).

### Quantifying habitat heterogeneity and urban index

We calculated habitat heterogeneity from the 3CS LC (Copernicus Climate Change Service Land Cover) and the ESA-CCI LC (European Space Agency-Climate Change Initiative Land Cover) land cover products (ESA. Land Cover CCI Product User Guide 2017; ESA. 3CS Land Cover Product User Guide 2020). These datasets provide methodologically consistent land cover per year and gridded maps from 1992 to 2019, with a global coverage and a spatial resolution of circa 300 m per pixel (0.002778° or 10 arcseconds). Each pixel in the products is classified as one of the 22 land cover categories defined by the UN-FAO-LCCS (United Nations Food and Agriculture Organization Land Cover Classification System). From a subset of studies included in our main meta-analysis, we could extract the coordinates of their urban and non-urban populations (26 studies out of 68 provided accurate coordinates of their urban and non-urban populations). Then, we sampled the landscape of every study by extracting the number of pixels belonging to each land cover category around each urban and non-urban location (i.e., within a circular buffer around each location). The extraction was performed for several buffer radii from 250 m to 5000 m in intervals of 250 m. Landscape heterogeneity was calculated as the effective number of land covers present in each buffer and computed as the exponential of the Shannon-Wiener diversity index (i.e., Hill’s numbers for *q* = 1) (Hill 1973; Chao *et al*. 2014), resulting into a measure that not only accounts for the absolute richness of land cover categories but also weights in the relative abundance of each category. An urban index was calculated as the proportion of each buffer area categorized as an ‘urban’ land cover type. Land cover data were processed and analysed using R (v.4.2.0; R Core Team 2022). Geospatial vectorial operations were conducted utilising the ‘sf’ R package (v.1.0-7; Pebesma 2018) while raster extractions were performed with the ‘raster’ R package (v.3.5-15; Hijmans 2020). All geospatial analyses were performed in the WSG 1984 projected Coordinate Reference Systems, EPSG: 6326. Additionally, we calculated the distance between each urban and non-urban pair of populations using the function ‘*pointDistance*’ in the R package ‘raster’. We could retrieve location information for 232 urban *versus* non-urban comparisons for laying date, clutch size and number of fledglings, from 11 species and 26 studies between 1992 and 2017 (land cover data were not available before 1992; see above).

### Meta-analyses

We handled the datasets, ran all analyses and produced visualisations using R (v.4.2.0; R Core Team 2022). To evaluate the effect of urbanisation on bird life-history traits, we fitted phylogenetic multilevel (intercept-only) meta-analyses for each response term (i.e., lnRR [Model 1] and lnCVR [Model 3]; Table 1) combining the three life-history traits (i.e., laying date, clutch size and number of fledglings; we also fitted models that separated variation between these traits; see below; Table 1). Both meta-analytic models estimated four random intercept effects, publication identity (i.e., among-study variation), population identity (i.e., in several cases, we found multiple studies from the same urban - non-urban populations pairs), phylogeny (more details below), species identity (i.e., among-species variation not explained by phylogeny), and an observation ID term. For the intercept-only models, we estimated total heterogeneity (*I*^2^) following Nakagawa & Santos (2012) and Senior et al. (2016b) as implemented in the R function ‘*i2_ml’* (‘orchaRd’ R package v.0.0.0.9000; Nakagawa et al. 2021).

**Table 1.** Description of meta-models. Model IDs are given sequentially from 1 to 10 to facilitate understanding of methods and results. ‘Data’ refers to whether a given model contained data for all traits of interest (‘All traits’) or models were fitted per trait. Moderator ‘Trait’ is a 3-level factor with levels ‘Laying date’, ‘Clutch size’ and ‘Number of fledglings’. ‘Equations’ provide references to the Equations described in the methods section, whereas ‘Details’ gives a brief description of each model Model ID and references to output tables and figures.

| Model ID | Response | Data | Moderators | Equations | Details |
| --- | --- | --- | --- | --- | --- |
| 1 | lnRR | All traits | Intercept | - | Overall meta-analysis. Univariate. Table S1. Figure S3. |
| 2 | lnRR | All traits | Trait | Equation 1 | Effect per trait. Trivariate. Tables S2, S3. Figure 2, 3, S4. |
| 3 | lnCVR | All traits | Intercept | - | Overall meta-analysis Univariate. Table S1. Figure S5. |
| 4 | lnCVR | All traits | Trait | Equation 1 | Effects per trait. Trivariate. Table 2, S4, S5. Figure 2, 3, S4. |
| 5 | lnCVR | All traits | Trait | Equation 1 | Comparison of intra-annual phenotypic variation. Table 2. |
| 6 | lnCVR | All traits | Trait | Equation 1, 7, 8 | Comparison of inter-annual phenotypic. Table 2. |
| 7 | lnCVR | All traits | Trait + Difference in urbanisation + Difference in habitat heterogeneity | Equation 1 (with additional moderators) | Trivariate. Fitted for different spatial scales. Figure 4. |
| 8 | SDHM | All traits | Trait | Equation 1 | Similar structure as Model 2. |
| 9 | lnVR | All traits | Trait | Equation 1 | Similar structure as Model 4. |
| 10 | lnSD | Each trait individually | Intercept + Habitat + lnMean | Equation 9 | Armed-based model (Senior *et al.* 2016a). |

### Phylogenies

Phylogenetic trees were extracted from the Open Tree of Life (Hinchliff *et al*. 2015; Rees & Cranston 2017), using the interface provided by the R package ‘rotl’ (v3.0.12; Michonneau *et al*. 2016; OpenTreeOfLife *et al*. 2019). We calculated tree branch length (Grafen 1989) and generated a phylogenetic correlation matrix to include in all our phylogenetic multilevel meta-analytic models (Figure 1). We assessed the phylogenetic signal in our meta-analysis based on the proportion of variation explained by the phylogeny (*I*^2^_phylogeny_; Cinar et al. 2022).

### Modelling heterogeneous variances and correlations among traits

Laying date, clutch size and number of fledglings are often correlated in bird species (Rowe *et al*. 1994; Dunn & Møller 2014). To assess whether urbanisation is associated with correlated responses across life-history traits and to test the robustness of our results to the existence of these correlations, we built trivariate meta-analytic models of lnRR and lnCVR that allowed us to simultaneously estimate trait-specific means (i.e., one intercept for each trait – Equation 1), trait-specific observation ID variances (i.e., one observation ID variance for each trait – Equation 1 & Equation 2) and trait-specific among-study variances and correlation among traits (Equation 1 & Equation 3). Including the random-effects detailed above, our model with heterogeneous variances and among-study correlations among traits can be written as: (we have omitted the term associated with sampling variance for simplicity – see Nakagawa et al. (2015) for more details)

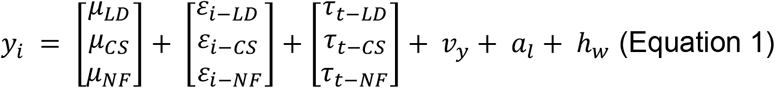

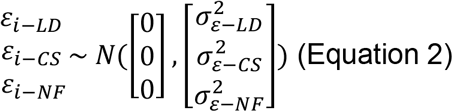

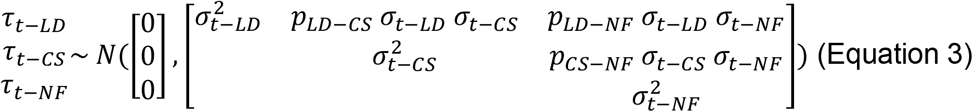

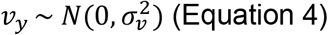

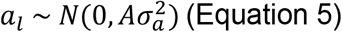

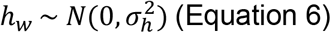

Where *y*_*i*_ is the statistic of interest (lnRR or lnCVR) for the *i*th effect size (*i* = 1,2,3, …, *k*; where *k* is the number of the effect sizes included in the analysis - i.e., number of urban – non-urban paired comparisons). ‘LD’, ‘CS’, ‘NF’ refer to overall means (μ), variances (σ^2^) and correlations (ρ) involving effect sizes for laying date (‘LD’), clutch size (‘CS’) and number of fledglings (‘NF’). *ε*_*i*_ is the observation ID deviation for the *i*th observation, which is assumed to follow a normal distribution with mean zero and variance 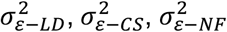 for laying date, clutch size and number of fledglings, respectively. *τ*_*t*−*LD*_, *τ*_*t*−*CS*_, and *τ*_*t*−*NF*_ are the deviations from the mean associated with the *t*th study and trait (‘LD’, ‘CS’, or ‘NF’), each following a multivariate normal distribution with mean of zero and variance-covariance structure detailed in Equation 5 (*p* provides the correlations between *τ*_*t*−*LD*_, *τ*_*t*−*CS*_, and *τ*_*t*−*NF*_). *v*_*y*_ provides the deviation from the overall mean associated with the *y*th population (Equation 4). *a*_*l*_ is the phylogenetic effect for the *l*th species, which follows a normal distribution with mean equal to zero and variance-covariance structure given by 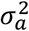, the variance of the phylogenetic effect, and *A*, a *l* by *l* matrix of distances between species calculated from a phylogenetic tree (Equation 5; details above). *h*_*w*_ captures among species variation not explained by the phylogenetic effect and follows a normal distribution around zero and variance 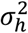 (Equation 6).

We compared models with different constraints in the parameters of the variance-covariance structure in Equation 3 to assess the strength of evidence for heterogeneous variances and correlations among traits (see results in Tables S2 and S4). We fitted these trivariate meta-analytic models in the ‘metafor’ R package (‘*rma*.*mv’* function; v3.4.0; Viechtbauer 2010) using maximum likelihood and compared models using AIC (Akaike Information Criterion; Burnham et al. 2011). We then calculated a ΔAIC value for each model (i.e., the difference in AIC between a given model and the model with the lowest AIC) and used this value to assess the strength of evidence for a given variance-covariance structure. We fitted models with the following constraints in the variance-covariance structure:

i. Single variance across traits and zero covariances:

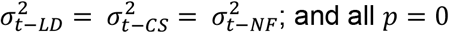
ii. Compound symmetric variance-covariance matrix:

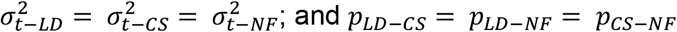
iii. Heteroscedastic compound symmetric variance-covariance matrix:

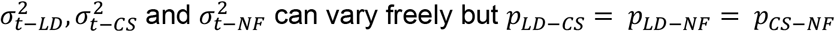
iv. Diagonal variance-covariance matrix:

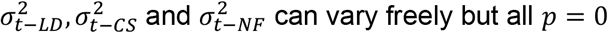
v. Unstructured variance-covariance matrix

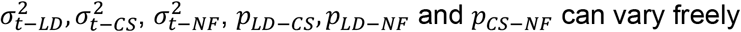

### Within- and between-breeding season differences in phenology and life-history traits

Urban and non-urban populations may differ in both within- and between-breeding season variation in life-history traits. However, differences in variation for these two temporal scales are likely generated by contrasting ecological and evolutionary processes. To disentangle processes operating at these two temporal scales, we performed additional meta-analyses including i) urban – non-urban comparisons within breeding seasons (k = 363 comparisons from 47 studies in the original dataset with effect sizes per year; Model 5) and ii) urban – non-urban comparisons between breeding seasons (i.e., combining all within-breeding season estimates from a study; k = 36 comparisons present in the original data set, plus 67 additional comparison calculated from within-breeding season estimates; see below). When a given study reported estimates across multiple breeding seasons, we calculated between-breeding season mean and variance as:

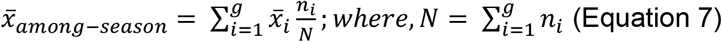

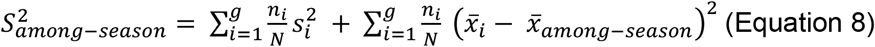

Where, 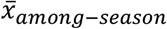 and 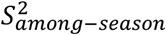 are mean and variance across multiple breeding seasons. *g* is the total number of breeding seasons reported by a given study; 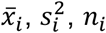, are mean, variance and sample size for each breeding season *i*. 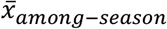 for a given study is, therefore, the weighted average across breeding seasons (Equation 7); whereas 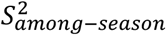 for a given study is the weighted sum of within-season variances (first term in Equation 8) and between-season mean variances (second term in Equation 8).

### Assessing the effect of urbanisation and habitat heterogeneity on differences in phenotypic variation between habitats

We investigated the spatial drivers of differences in phenotypic variation between urban and non-urban populations using the subset of studies which allowed the quantification of an urban index in urban and non-urban populations (see above). We first verified that the urban index was indeed higher for urban than for non-urban populations. We compared the urban index in urban and non-urban populations at different spatial scales via linear models, with the difference in urban index between population as the response variable and an intercept term. Then, to assess whether the increase in phenotypic variation in urban habitats was predicted by habitat heterogeneity and/or urban index, we ran an additional meta-regression to explain differences in phenotypic variation between urban and non-urban populations (i.e., lnCVR), where the difference in habitat heterogeneity and urban index between urban and non-urban populations were included as continuous moderators. This meta-regression included 232 urban – non-urban comparisons from 11 species and 26 studies (i.e., the subset of observations after 1992 for which geolocations were available).

### Sensitivity analyses

We assessed the robustness of our results with several complementary analyses. First, we re-ran the trivariate lnRR model (Model 2; Table 1) using Hedges’ g (Hedges 1981) with heteroscedastic population variances as the response variable (Table 1; Model 8; i.e., ‘SMDH’, calculated using the R function ‘*escalc*’ in the ‘metafor’ R package (v3.4.0; Viechtbauer 2010)). In addition, we assessed the robustness of the lnCVR results by re-running the trivariate lnCVR model (Model 4; Table 1) using lnVR as the response variable (i.e., the logarithm of the total variation ration; Nakagawa et al. 2015; Model 9; Table 1). Last, we used an alternative approach that directly models the log of the phenotypic standard deviation (lnSD) to assess differences in phenotypic variation between urban and non-urban populations (Eq. 18 in Nakagawa et al. (2015); Model 10; Table 1). We followed the model specification shown in Senior et al. (2016a), in short:

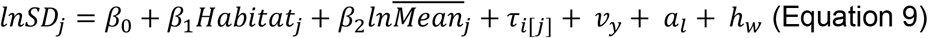

Where *β*_0_ is the overall intercept, *β*_1_ is the habitat effect on lnSD (i.e., a *β*_1_ statistically different from zero would indicate that urban and non-urban populations differ in their phenotypic variation) and *β*_2_ is the slope of the regression of (log) mean values against (log) standard deviations, which is explicitly modelled. *v*_*y*_, *a*_*l*_ and *h*_*w*_ are as per Equation 1. *τ*_*i*[*j*]_ is the random effect for the *j*th effect size in the *i*th study. Within each study effect sizes across habitats are assumed to be correlated; this correlation is calculated by the model (Senior *et al*. 2016a). We applied the model in Equation 9 for each trait independently (i.e., three univariate models, one per trait).

### Publication bias

We assessed the evidence for the existence of two types of publication biases, small-study and decline effects (time-lag effects), following Nakagawa et al. (2022). For that, we ran four additional uni-moderator multilevel meta-analytic models, two for lnRR and two lnCVR. Each of these models included as a single moderator either the square-root of the inverse of the effective sample size or the mean-centered year of study publication (Trikalinos & Ioannidis 2005; Nakagawa *et al*. 2022). The variation explained by these moderators (i.e., R^2^_marginal_) was calculated using the R function ‘r*2_ml’* (‘orchaRd’ R package v.0.0.0.9000; Nakagawa et al. 2021).

## Results

After systematically inspecting 1,166 studies published between 1958 and 2020 (Figure S1), our meta-analysis included 399 urban – non-urban comparisons for three bird life-history traits: laying date (k = 151 effect sizes, n = 32 studies), clutch size (k = 119 effect sizes, n = 42 studies) and number of fledglings (k = 129 effect sizes, n = 48 studies) (Figure 1). This dataset included 35 bird species, with most studies located in the northern hemisphere (Figure 1c).

### i) Is urbanisation associated with shifts in mean life-history traits?

We found that urban populations tended to have, on average, 3.6% lower mean values than their non-urban counterparts, but note that the 95% confidence interval (hereafter ‘CI’) for this estimate overlapped zero (Model 1: lnRR mean estimate [95% CI] = −0.035 [−0.076,0.005]; Figure S3; Table S1). Total heterogeneity was high (*I*^2^_total_ = 97.8%), with 17.2% of it explained by phylogenetic and species-specific effects combined (*I*^2^_phylogeny_ = 1.7%; *I*^2^_study ID_ = 15.5%), while 8.4% was explained by differences among studies (Table S1). Further analyses calculating urban effects per trait and accounting for potential covariation in the response to urbanisation across the three focal traits (i.e., using a model with an unstructured variance-covariance matrix; see Methods and Table S2) confirmed that urban populations had indeed lower mean values in every life-history trait: urban populations laid their eggs earlier (Model 2: lnRR [95% CI] = −0.048 [−0.084, −0.012]; Figure 2a), laid smaller clutches (Model 2: lnRR [95% CI] = −0.066 [−0.107, −0.025]; Figure 2a), and tended to produce fewer fledglings per clutch than non-urban populations (Model 2: lnRR [95% CI] = −0.070 [−0.171, 0.032]; Figure 2a). This meta-analytic model estimated different random effect intercepts per trait and allowed for correlations across traits (Model 2; see Methods for details). This model revealed correlations in the response to urbanisation across traits: studies reporting earlier laying date in urban populations also reported more similar clutch size and number of fledglings between populations (i.e., negative correlations between lnRR for laying dates and clutch size; Figure 3a & 3b). Likewise, studies reporting large differences in clutch size between urban and non-urban populations also reported large differences between both habitats in number of fledglings (Figure 3c; see ‘Study ID (correlations)’ in Table S3; i.e., correlations among studies in the values of lnRR for each trait).

**Figure 2.**
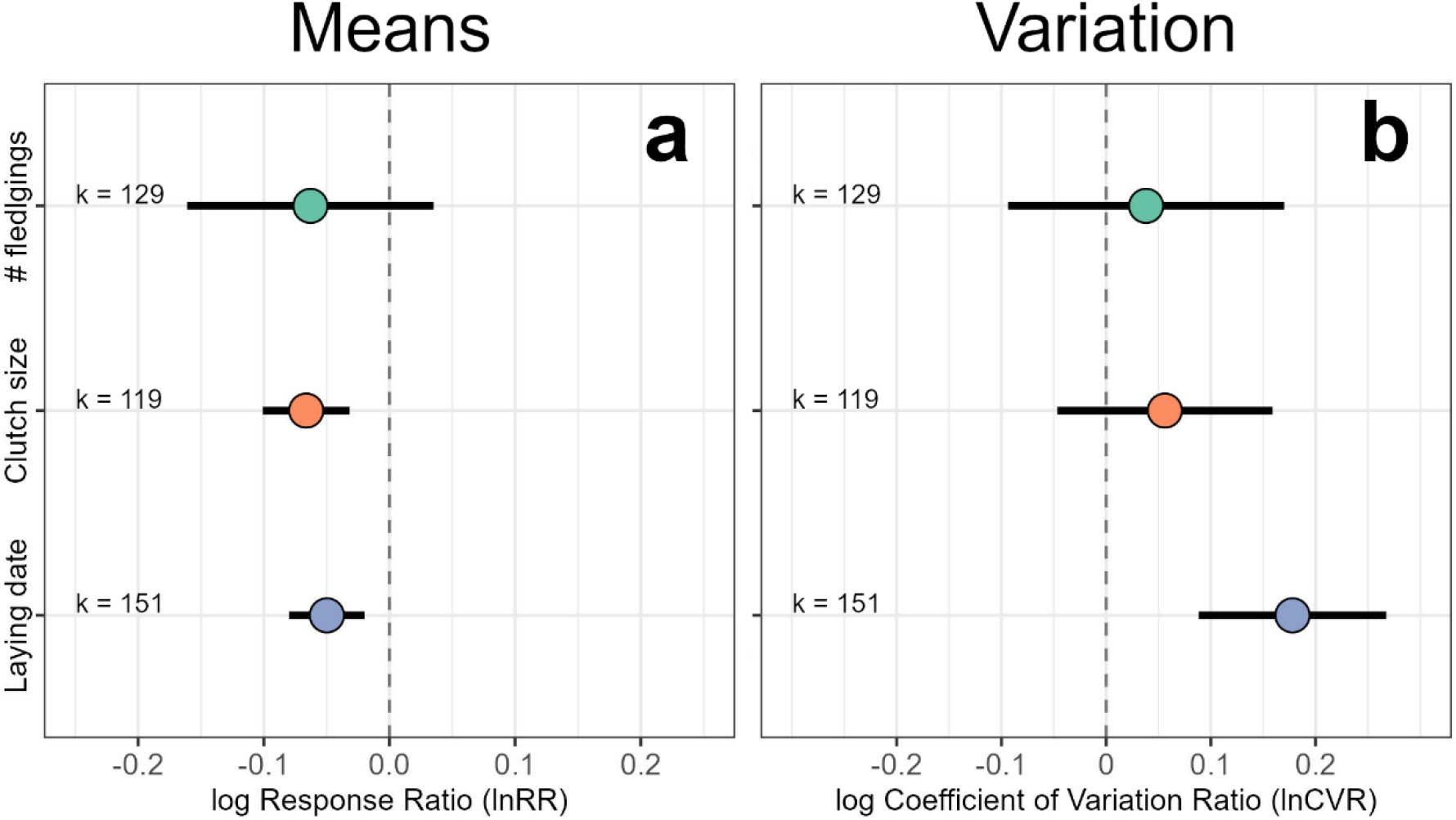
Urban populations have earlier phenology, lower reproductive output and more variable life-history traits than non-urban populations. (**a**) Urban populations laid earlier and had smaller clutches, producing fewer fledglings, than their paired non-urban populations (illustrated by negative lnRR estimates; Model 2). (**b**) Our meta-analysis revealed that variation in life-history traits was higher in urban populations compared to non-urban counterparts, with a marked difference between populations in laying date (illustrated by positive estimates of lnCVR; Model 4). Model estimates for (**a**) lnRR and (**b**) lnCVR are shown along with their 95% confidence intervals per trait as calculated by our phylogenetic multilevel meta-analytic models accounting for correlated responses to urbanisation among traits (see Table S3 & Table S5 for full model outputs and Figure S3 and S5 for overall meta-analyses of lnRR and lnCVR). Raw data and model estimates are presented in Figure S4. ‘k’ provides the number of urban – non-urban comparisons.

**Figure 3.**
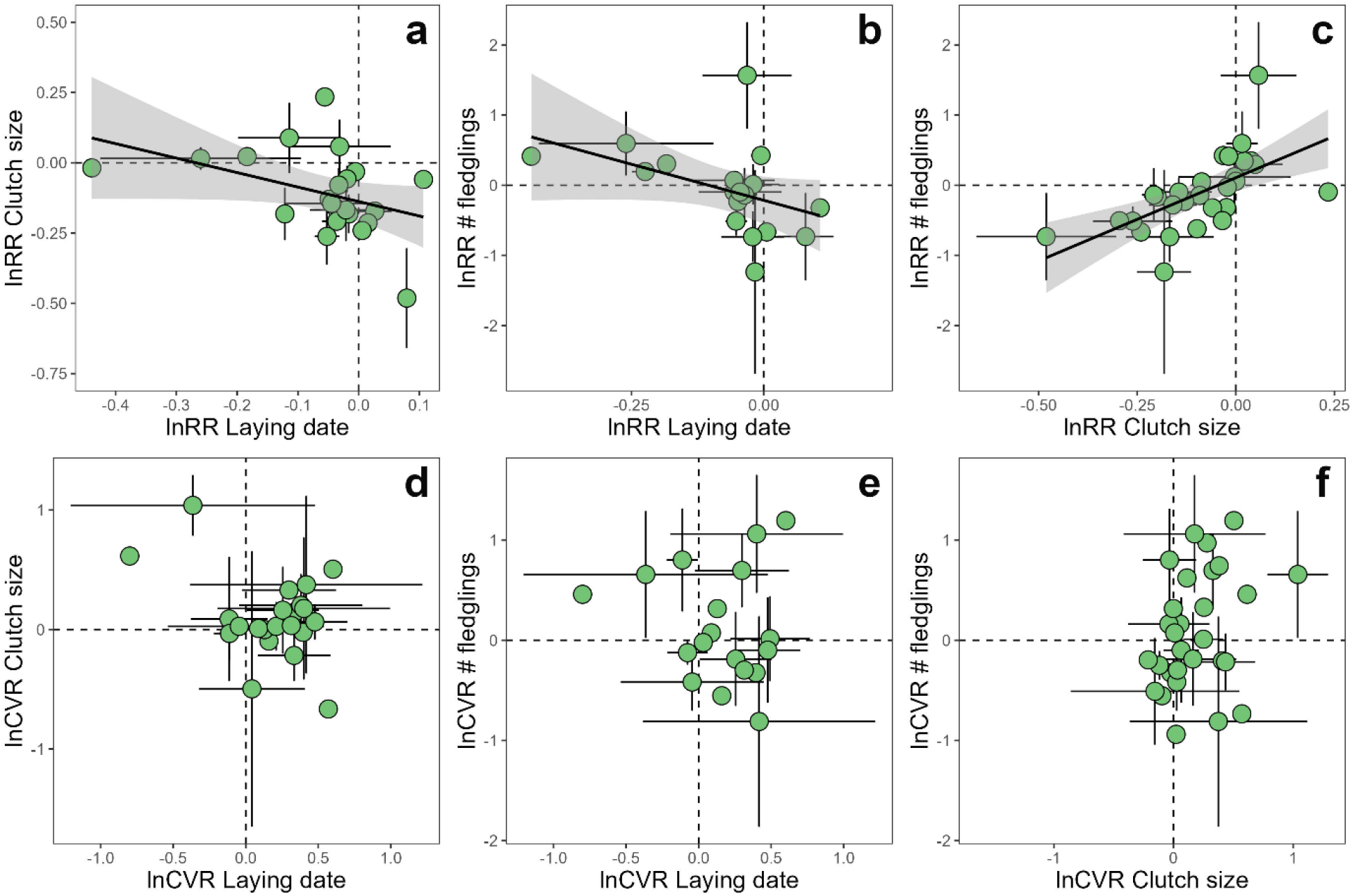
Life-history traits show a correlated response to urbanisation. Our meta-analysis investigated correlated responses to urbanisation across the three studied life-history traits, and revealed strong correlations in log response ratio (lnRR) but not log coefficient of variation ratio (lnCVR). (**a**) Earlier laying dates in urban populations compared to non-urban counterparts (i.e., negative values in the x axis) were associated with no differences in clutch size across habitats (i.e., y axis values close to zero), leading to a negative correlation between lnRR for these two traits. (**b**) A similar pattern was found between lnRR for laying dates and number of fledglings, while (**c**) lnRR for clutch size and number of fledglings were positively correlated (Table S2; Table S3; Model 2). (**d** - **f**) We found no strong statistical evidence for models including correlations across traits in how urbanisation affected phenotypic variation (Table S4, Table S5): (**d**) differences between habitats in phenotypic variation in laying dates were not associated with differences between habitats in phenotypic variation in clutch size or (**e**) number of fledglings; and (**f**) differences between habitats in variation in clutch size were not associated with differences between habitats in variation in number of fledglings. Points represent mean raw values per study ± SE. Regression lines (mean ± SE) in **a** - **c** were fitted using linear regressions to illustrate the correlation revealed by our trivariate meta-analysis (Model 2; Table S3).

### ii) Is urbanisation associated with changes in variation in life-history traits?

The coefficient of phenotypic variation in urban populations was, on average, 4.4% higher than in non-urban populations, but note that the 95%CI for this estimate overlapped zero (Model 3: lnCVR mean estimate [95% CI] = 0.043 [−0.092, 0.178]; *I*^2^_total_ = 74.3%; Figure S5 and Table S1). 9.1% of the heterogeneity in lnCVR was explained by phylogenetic and species-specific effects combined (*I*^2^_phylogeny_ = 5.8%; *I*^2^_species ID_ = 3.3%), while differences between studies explained no heterogeneity in lnCVR (*I*^2^_species ID_ = 0.0%; Table S1). A subsequent model of lnCVR separating urban effects on phenotypic variation per trait and accounting for potential covariation across the three investigated traits in the response to urbanisation (see Methods and Table S4) revealed that the overall effect of urbanisation on life-history trait variation was driven by urban populations having a more variable phenology than their non-urban counterparts (Model 4: lnCVR mean for laying date [95% CI] = 0.176 [0.084, 0.268], i.e., 19.2% more variation, on average, in laying date in urban than non-urban populations). Although the 95%CIs overlapped zero, the direction of the average effects for clutch size and number of fledglings also reflected higher phenotypic variation in urban compared to non-urban populations (Model 4: lnCVR mean estimates [95% CI]: clutch size = 0.055 [−0.051, 0.160], number of fledglings = 0.037 [−0.096, 0.171]; Figure 2b). We did not find evidence for correlations in lnCVR between the three life-history traits (Figure 3; the model including correlations among traits scored more than 1.08 AIC points below the top model, which only included independent Study ID random intercepts per trait [Model 4]; Table S4; Table S5).

### iii) What is the temporal and spatial scale at which urbanisation affects phenotypic variation?

Differences in phenotypic variation in laying date between the urban and non-urban populations arose from differences in variation within breeding seasons (i.e., intra-annual) rather than between breeding seasons (i.e., inter-annual; Table 2). While laying dates in urban populations were more variable than in non-urban populations within breeding seasons (Model 5: lnCVR mean estimate [95% CI] = 0.177 [0.078, 0.281]; Table 2), a subsequent meta-analytic model isolating effects on phenotypic variation arising from between breeding season fluctuations revealed no difference between urban and non-urban populations (Model 6: lnCVR intercept mean [95% CI] = 0.074 [−0.014, 0.161]; Table 2). The sample size for this latter meta-analysis was almost four times smaller than for the meta-analysis of within breeding season differences in variation; however, the lnCVR estimates were very different between these models: the mean lnCVR within breeding seasons was more than 2.4 times larger than the mean lnCVR among breeding seasons (Table 2).

**Table 2.** Differences in variation (lnCVR) in life-history traits between urban and non-urban populations at different temporal scales. Urban – non-urban differences in variation (lnCVR) in laying date, clutch size and number of fledglings per clutch were meta-analysed to assess differences in variation between urban and non-urban populations within (‘intra-annual’) and among (‘inter-annual’) breeding seasons (e.g., different temporal scales). lnCVR estimates represent meta-analytic model intercepts following the model structure presented in Table S5; positive values indicate higher variation in urban populations than in non-urban populations and *vice versa*. ‘CI’ = confidence interval; ‘k’ = sample size. Terms in italic bold highlight lnCVR estimates whose 95%CIs do not overlap zero. See Table 1 for a description of model IDs.

| Temporal scale | lnCVR estimate [95% CI] |  |  | k |
| --- | --- | --- | --- | --- |
|  | Laying date | Clutch size | Number of fledglings |  |
| **Overall**<br>**[Model 4]** | ***0.176***<br>***[0.084, 0.268]*** | 0.055<br>[-0.051, 0.160] | 0.037<br>[-0.096, 0.171] | 399 |
| **Intra-annual**<br>**[Model 5]** | ***0.177***<br>***[0.078, 0.282]*** | 0.015<br>[-0.122, 0.152] | 0.116<br>[-0.059, 0.291] | 363 |
| **Inter-annual**<br>**[Model 6]** | 0.074<br>[-0.014, 0.161] | 0.096<br>[-0.019, 0.211] | -0.006<br>[-0.147, 0.135] | 103 |

Furthermore, to assess whether urbanisation and/or habitat heterogeneity could explain increased phenotypic variation in urban bird populations, we investigated the extent to which our quantification of urban index and habitat heterogeneity predicted differences in phenotypic variation across populations. First, we confirmed that the urban populations included in our meta-analysis showed higher levels of urbanisation than paired non-urban populations regardless of the spatial scale used (urban index in urban population ± SE = 0.669 ± 0.047; urban index in non-urban population ± SE = 0.021 ± 0.007; at a spatial scale of 2000 m in both cases for reference; Figure 4a). Including the difference in urban index and habitat heterogeneity between paired urban and non-urban populations as a moderator in a meta-regression revealed that the more heterogeneous the urban habitat was, the larger the phenotypic variation in this habitat compared to the non-urban habitat; this effect was particularly strong at medium-large spatial scales (Figure 4c). Differences in urban index between populations did not strongly explain variation in lnCVR (Figure 4b). Urban and non-urban populations in each studied pair were located at a mean distance of 65.4 km (median = 33.1 km; range = [2.4 km, 625.1 km]; n = 26 geo-referenced studies).

**Figure 4.**
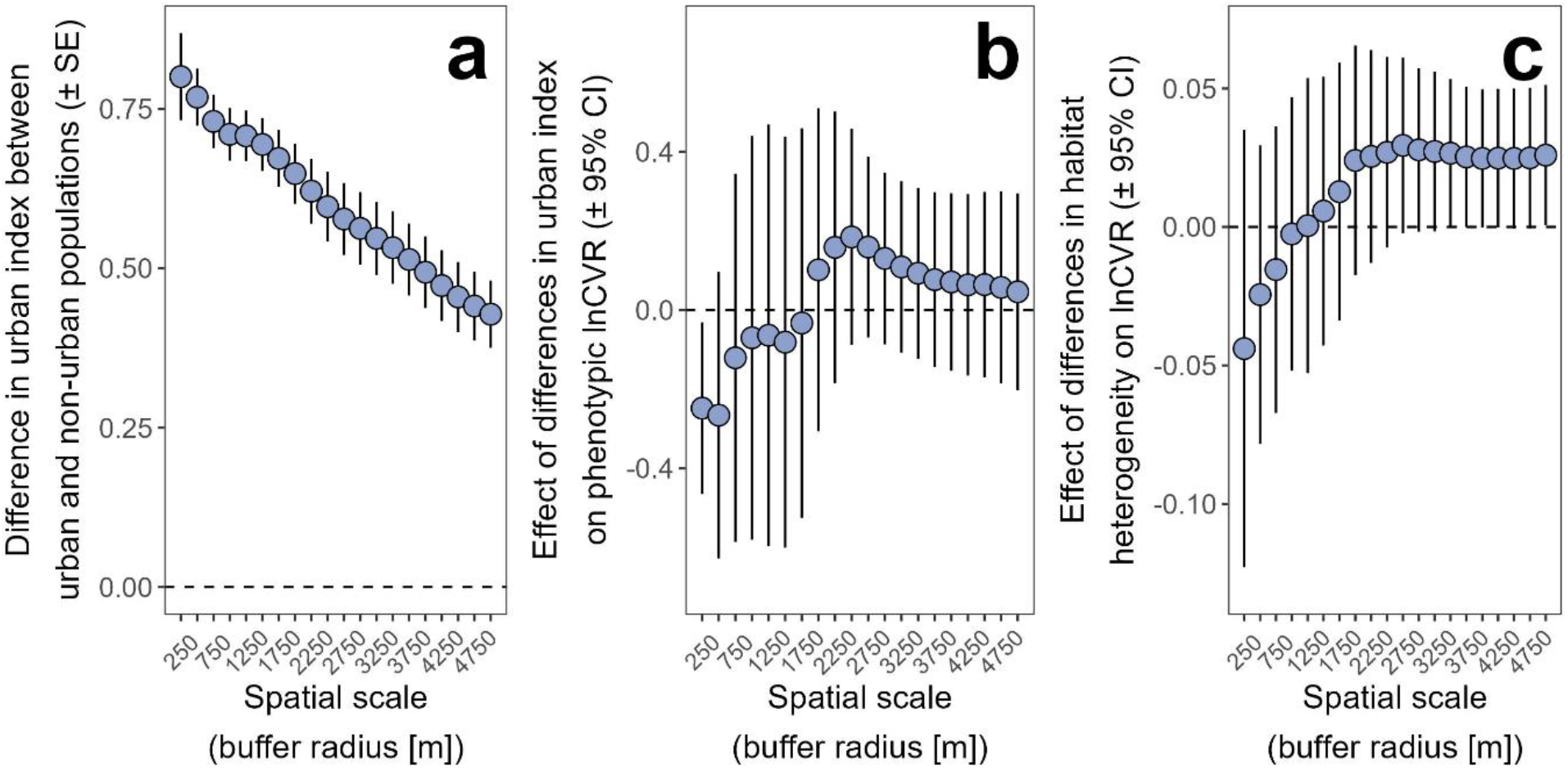
Effects of habitat heterogeneity on the difference in phenotypic variation between urban and non-urban bird populations (i.e., lnCVR). (**a**) After quantifying urban index and habitat heterogeneity, we verified that urban populations had higher urban index (i.e., the proportion of landcover at a given spatial scale categorised as ‘urban’ [see methods]). The y axis represents the difference in urban index between urban and non-urban populations. The positive values observed for all comparisons represent that urban populations had higher urban index than their non-urban neighbours. (**b**) Differences in urban index between urban and non-urban populations did not predict the magnitude of the difference in phenotypic variation between populations (i.e., lnCVR). This figure shows the estimated effect of differences in urban index between populations on lnCVR. Positive values indicate that the higher the difference in urban index between urban and non-urban populations, the higher the lnCVR value (i.e., larger values of phenotypic variation in urban populations compared to non-urban counterparts). (**c**) Differences in habitat heterogeneity between urban and non-urban populations did positively predict the magnitude of the difference in phenotypic variation between populations (i.e., lnCVR), particularly at large spatial scales. This figure shows the estimated effect of differences in habitat heterogeneity on lnCVR at different spatial scales. Positive values indicate that the higher the difference in habitat heterogeneity between urban and non-urban populations, the higher the lnCVR value (i.e., larger values of phenotypic variation in urban populations compared to non-urban counterparts). Points represent mean model estimates ± SE in **a**, and mean model estimates ± 95% confidence intervals (95%CI) in **b** and **c**. ‘Spatial scale’ refers to the radius of a circular area centred at each study location and over which urban index and habitat heterogeneity was calculated.

### iv) Sensitivity analyses and assessment of publication bias

In line with our main analysis of lnRR (Table S3), using SMDH as the effect size provided negative estimates (i.e., lower phenotypic means in urban populations) for laying dates (SMDH mean estimate [95% CI] = −0.298 [−0.634, 0.039]), clutch size (SMDH mean estimate [95% CI] = −0.145 [−0.420, 0.130]) and number of fledglings (SMDH mean estimate [95% CI] = −0.022 [−0.298, 0.254]) (Model 8 in Table 1). Uncertainty around mean SMDH estimates was high and the 95%Cis overlapped zero. Analysing lnVR instead of lnCVR provided further evidence for increased phenotypic variation in urban populations, particularly for phenology (Model 9 in Table 1): the mean lnVR estimate for laying date was positive and statistically different from zero (lnVR mean estimate for laying date [95% CI] = 0.158 [0.069, 0.247]). As in our lnCVR, lnVR mean estimates for clutch size and number of fledglings were close to zero (lnVR mean estimate for clutch size [95% CI] = −0.012 [−0.110, 0.056]; lnVR mean estimate for number of fledglings [95% CI] = −0.034 [−0.120, 0.052]). Additionally, the arm-based model of lnSD for laying date (Model 10 in Table 1) revealed a positive ‘urban’ effect on lnSD: urban populations had lnSD values 0.197 higher than non-urban populations (i.e., *β*_1_ in Equation 9; 95%CI = [0.122, 0.272]). Laying date (log) mean phenotypic values were positively correlated with lnSD (i.e., *β*_2_ in Equation 9; estimate [95%CI] = 0.416 [0.068, 0.764]). The arm-based models of clutch size and number of fledglings confirmed correlations between lnMean and lnSD (*β*_2_ in Equation 9 for clutch size, estimate [95%CI] = 0.326 [0.070, 0.582]; for number of fledglings, estimate [95%CI] = 0.231 [0.155, 0.307]), but did not provide evidence for urban effects on phenotypic variation in clutch size or number of fledglings (*β*_1_ in Equation 9 for clutch size, estimate [95%CI] = 0.020 [−0.079, 0.119]; for number of fledglings, estimate [95%CI] = −0.017 [−0.099, 0.065]). We did not find evidence of publication bias in lnRR or lnCVR (Supplementary Text C).

## Discussion

We compiled a global dataset of bird life-history traits for paired urban and non-urban populations of the same species to assess how urban living is related to changes in phenotypic means and variation for breeding phenology, reproductive effort, and reproductive success. A phylogenetically controlled multilevel meta-analysis of this dataset confirms a well-documented effect of urbanisation on mean phenotypes: urban bird populations lay earlier and smaller clutches than their non-urban counterparts. This model, however, also reveals correlated responses to urbanisation across life-history traits: e.g., the earlier the laying date in urban populations, the smaller the difference in clutch sizes between habitats. Our study goes a step further than previous meta-analyses in urban ecology by explicitly investigating how urbanisation could impact phenotypic variation. Our findings highlight that urbanisation is associated with both a decrease in mean phenotypes, and an increase phenotypic variation. Investigating the temporal and spatial scale at which urban phenotypic variation increases revealed hints at the ecological causes and evolutionary consequences.

Urbanisation has been associated with shifts in mean phenotypic values across a many organisms (Alberti *et al*. 2017; Merckx *et al*. 2018; Santangelo *et al*. 2022), including birds, which generally show smaller body sizes and lower life-history trait values in urban habitats (Chamberlain *et al*. 2009; Sepp *et al*. 2018; Thompson *et al*. 2022). Our analyses expand the spatial, temporal and phylogenetic coverage of previous meta-analyses of the avian literature (Chamberlain *et al*. 2009; Sepp *et al*. 2018), and agree on their findings. Our results indicate that urban bird populations lay their eggs earlier and produce smaller clutches, which results in a lower number of surviving nestlings, than their non-urban neighbouring populations. Note, that our analysis indicates a high total heterogeneity in lnRR (*I*^2^_total_ = 97.8%). This finding indicates large variation (e.g., among studies and species) in how urbanisation associates with changes in mean phenotypes and suggests that additional ecological traits (e.g., diet or migratory strategy) may also affect how populations respond to urbanisation. Our results also indicate that the mean response to urbanisation is correlated among traits. Interestingly, we found that the earlier the laying dates were in urban *versus* non-urban populations, the smaller the difference in clutch size and in number of surviving nestlings between habitats. Many bird species show a negative phenotypic and genetic correlation between clutch size and lay date (Rowe *et al*. 1994; Sheldon *et al*. 2003; Dunn & Møller 2014), and these two traits are often hypothesized to co-evolve (Garant *et al*. 2008). All else being equal, urban conditions triggering an earlier onset of reproduction (because of e.g., light pollution (Dominoni *et al*. 2013) or increased resource availability during winter (Schoech *et al*. 2004)) could indirectly increase clutch size and, therefore, reduce differences in reproductive output between urban and non-urban populations that arise via other mechanisms (e.g., resource limitation in spring; Seress *et al*. 2018, 2020). The extent to which mean phenotypic shifts represent adaptive responses to urbanisation in birds, either via genetic changes or plasticity, or are maladaptive, is mostly unknown (Lambert *et al*. 2020; Branston *et al*. 2021; Caizergues *et al*. 2022; Santangelo *et al*. 2022). Our results, however, highlight that phenotypic shifts in urban populations are widespread and that the response to urbanisation of associated life-history traits should be investigated together.

Urbanisation has been recently hypothesised to increase phenotypic variation and, indeed, higher variation in morphological traits of urban great tits (*Parus major*) and blue tits (*Cyanistes caeruleus*) has been recently reported (Thompson *et al*. 2022). Our findings greatly expand the evidence for this emerging hypothesis showing that urbanisation is overall associated with increases in variation in laying date across many bird species. Previous studies have suggested that city characteristics, such as warmer temperatures in early spring due to the urban heat island effect, could allow birds to lay more clutches per season (Yeh & Price 2004; Schoech *et al*. 2008), with thereby longer breeding seasons and hence higher phenotypic variation in urban laying dates (a similar result has also been reported in Lepidoptera; Merckx *et al*. 2021). This effect, however, does not necessarily explain our results as our meta-analysis only included first clutch laying dates per season. As such, our findings indicate that urban bird populations display more variation in the *onset* of reproduction than their non-urban neighbours.

Higher phenotypic variation in urban than in non-urban populations within breeding seasons could be explained by at least two, non-exclusive, eco-evolutionary mechanisms: differences in the underlying additive genetic variance in laying date, whereby urban birds have a wider range of breeding values for laying date; and / or differences in habitat heterogeneity influencing plasticity in laying date, whereby urban areas have larger environmental variation than non-urban habitats (Shochat *et al*. 2006; Heisler & Brazel 2018; Strubbe *et al*. 2020; Thompson *et al*. 2022). No study to date has investigated whether urban birds show higher additive genetic variance than non-urban populations. However, genetic analyses of European great tits in urban and non-urban habitats generally suggest small differences in the magnitude of genetic variation between habitats (Björklund et al. 2010; Caizergues et al. 2021; Salmón et al. 2021). This is, perhaps, not surprising given the high mobility of birds and the fact that gene flow between urban and non-urban bird populations likely occurs at a large spatial scale (Salmón *et al*. 2021). Interestingly, some studies have reported weaker selection for laying date in urban areas than in non-urban habitats, suggesting relaxed selection on phenology in urban birds (Caizergues *et al*. 2018; Branston *et al*. 2021), which could increase genetic variation in phenology. Assessing differences in phenotypic variation between urban and non-urban populations of less mobile species will be important to evaluate how biological traits (e.g., dispersal ability) determine the evolutionary impact of urban ecological conditions. To this end, previous work in mammal and amphibian species that have a lower dispersal ability than birds suggests a similar level of (genetic) variation between urban and non-urban habitats (Fusco *et al*. 2021; Richardson *et al*. 2021).

Habitat complexity differs between urban and non-urban habitats (Arnfield 2003; Pickett *et al*. 2017). Our analyses indicate that differences in urban *versus* non-urban habitat heterogeneity could indeed help to explain the observed pattern of increased phenotypic variation in urban populations. Several ecological mechanisms could mediate this effect. Urban environments are characterised by an array of microhabitats with varying levels of human pressure, exotic plant species and resource availability. Thus, the intensity and timing of the environmental cues that birds use to time their reproduction could vary at a small local scale, increasing phenotypic variation in phenology in the presence of plasticity. The existence of plastic responses to urban habitat heterogeneity, which our results might indicate, do not preclude selection from acting on urban bird populations. First, plasticity is an important mechanism of adaptation, sometimes aligned in direction with adaptative genetic changes (De Lisle *et al*. 2022), and indeed is often involved in adaptation to urban environments (Halfwerk *et al*. 2019; Campbell-Staton *et al*. 2021). Second, plastic responses can aid adaptation to urban conditions in the presence of genetic-by-environment interactions by increasing genetic variation available for natural selection (Via & Lande 1985). Addressing which evolutionary mechanisms cause the observed increase in phenotypic variation in urban bird populations is beyond the scope of this study and we acknowledge that these arguments are largely speculative at this point. However, our findings highlight that eco-evolutionary processes could largely differ between urban and non-urban bird populations and generate new avenues for future research in urban ecology and evolution.

In agreement with our initial predictions, habitat heterogeneity was associated with the magnitude of the difference in phenotypic variation between urban and non-urban bird populations. However, we acknowledge that this analysis has several limitations and that the results require cautious interpretation. First, only a subset of published studies provided coordinates for their urban and non-urban study populations (30 out of 65 published papers). When study site coordinates were provided, only one pair of coordinates per study location was provided, preventing an accurate assessment of the actual area over which a given breeding population was studied. Additionally, it is common in urban eco-evolutionary studies to monitor several populations within one single city. However, in most studies, spatial information was provided at the scale of the whole city (e.g., a single set of coordinates), preventing the accurate quantification of habitat heterogeneity for every sub-population within a given urban habitat. These limitations highlight that the ability to perform global meta-analyses on the effects of urban habitat heterogeneity on phenotypic variation would be greatly improved if individual studies in urban ecology provided accurate coordinates of the location of their study populations. Reporting such information would allow future research synthesis to quantify phenotypic variation within urban populations (e.g., across different sub-populations in the same city) and between urban and non-urban populations.

Taken together, our results show that urbanisation is associated with both a decrease in mean phenotypic values and increasing phenotypic variation in bird populations. Our analyses also highlight a temporal and spatial mechanism that could generate such differences in phenotypic variation between urban and non-urban habitats. We show that urban bird populations have a more variable phenology than non-urban conspecifics within breeding seasons (i.e., differences in phenology across habitats are seemingly not due to between-year fluctuations) suggesting that the ecological conditions that generate such differences are constant across multiple years. Our coupled spatial analysis indicates habitat heterogeneity and plastic responses as potential eco-evolutionary drivers generating these results. The eco-evolutionary implications of higher phenotypic variation in urban environments will likely vary among species (Thompson *et al*. 2022) and our findings highlight the need for detailed investigation of these consequences. To this end, long-term studies of individually marked organisms in replicated paired urban and non-urban environments could be particularly fruitful to unravel whether differences in phenotypic variation between urban and non-urban populations are caused by differences in underlying genetic variation and/or plastic responses to the urban environment.

## Acknowledgements

We would like to thank Paul Bellamy, Clint Boal, Peter Ferns, Michał Glądalski, Shelley A. Hinsley, Piotr Minias, Christy Morrissey, Matthew Reudink, Staffan Roos, Renaud Scheifler, Christine Stracey, Daniel Shustack and Jarosław Wawrzyniak for kindly replying to our data queries. We are grateful to Antje Girndt for developing the list of avian genera included in our search string.

## Funding

PC-L, CJB and DMD were funded by a Highlight Topics grant from the Natural Environment Research Council awarded to DMD (NE/S005773/1).

## Competing interests

The authors declare no competing interests.

## Notes

https://github.com/PabloCapilla/meta-analysis_variation_urban

### Competing Interest Statement

The authors have declared no competing interest.

### Summary of Updates

Revised version

## References

Alberti, M., Correa, C., Marzluff, J.M., Hendry, A.P., Palkovacs, E.P., Gotanda, K.M., et al. (2017). Global urban signatures of phenotypic change in animal and plant populations. Proceedings of the National Academy of Science, 114, 8951–8956.

Antonov, A. & Atanasova, D.Y. (2003). Small-scale differences in the breeding ecology of urban and rural Magpies Pica pica. Ornis Fennica, 80, 21–30.

Arnfield, A.J. (2003). Two decades of urban climate research: A review of turbulence, exchanges of energy and water, and the urban heat island. International Journal of Climatology, 23, 1–26.

Bailly, J., Scheifler, R., Berthe, S., Clément-Demange, V.A., Leblond, M., Pasteur, B., et al. (2016). From eggs to fledging: Negative impact of urban habitat on reproduction in two tit species. Journal of Ornithology, 157, 377–392.

Baldan, D. & Ouyang, J.Q. (2020). Urban resources limit pair coordination over offspring provisioning. Scientific Reports, 10, 15888.

Beck, N.R. & Heinsohn, R. (2006). Group composition and reproductive success of cooperatively breeding white-winged choughs (Corcorax melanorhamphos) in urban and non-urban habitat. Austral Ecology, 31, 588–596.

Berardelli, D., Desmond, M.J. & Murray, L. (2010). Reproductive success of burrowing owls in urban and grassland habitats in southern New Mexico. Wilson Journal of Ornithology, 122, 51–59.

Biard, C., Brischoux, F., Meillère, A., Michaud, B., Nivière, M., Ruault, S., et al. (2017). Growing in cities: An urban penalty for wild birds? A study of phenotypic differences between urban and rural great tit chicks (Parus major). Frontiers in Ecology and Evolution, 5, 79.

Björklund, M., Ruiz, I. & Senar, J.C. (2010). Genetic differentiation in the urban habitat: the great tits (Parus major) of the parks of Barcelona city. Biological Journal of the Linnean Society, 99, 9–19.

Boal, C.W. & Mannan, R.. W. (1999). Comparative Breeding Ecology of Cooper’s Hawks in Urban and Exurban Areas of Southeastern Arizona. The Journal of Wildlife Management, 63, 77–84.

Bobek, O., Gal, A., Saltz, D. & Motro, U. (2018). Effect of nest-site microclimatic conditions on nesting success in the Lesser Kestrel Falco naumanni. Bird Study, 65, 444–450.

Brahmia, Z., Scheifler, R., Crini, N., Maas, S., Giraudoux, P. & Benyacoub, S. (2013). Breeding performance of blue tits (Cyanistes cæruleus ultramarinus) in relation to lead pollution and nest failure rates in rural, intermediate, and urban sites in Algeria. Environmental Pollution, 174, 171–178.

Branston, C.J., Capilla, P., Christopher, L., Griffiths, K., White, S. & Dominoni, D.M. (2021). Urbanisation weakens selection on the timing of breeding and clutch size in blue tits but not in great tits. Behavioral Ecology and Sociobiology, 75, 155.

Burnham, K.P., Anderson, D.R. & Huyvaert, K.P. (2011). AIC model selection and multimodel inference in behavioral ecology: some background, observations, and comparisons. Behavioral Ecology and Sociobiology, 65, 23–35.

Caizergues, A.E., Charmantier, A. & Grégoire, A. (2018). Urban versus forest ecotypes are not explained by divergent reproductive selection. Proceedings of the Royal Society B, 285, 20180261.

Caizergues, A.E., Grégoire, A., Choquet, R., Perret, S. & Charmantier, A. (2022). Are behaviour and stress-related phenotypes in urban birds adaptive? Journal of Animal Ecology, 00, 1–15.

Caizergues, A.E., le Luyer, J., Grégoire, A., Szulkin, M., Senar, J., Charmantier, A., et al. (2021). Epigenetics and the city: non-parallel DNA methylation modifications across pairs of urban-rural Great tit populations. Evolutionary Applications, 15, 149–165.

Campbell-Staton, S.C., Velotta, J.P. & Winchell, K.M. (2021). Selection on adaptive and maladaptive gene expression plasticity during thermal adaptation to urban heat islands. Nature Communications, 12, 6195.

Campbell-Staton, S.C., Winchell, K.M., Rochette, N.C., Fredette, J., Maayan, I., Schweizer, R.M., et al. (2020). Parallel selection on thermal physiology facilitates repeated adaptation of city lizards to urban heat islands. Nature Ecology & Evolution, 4, 652–658.

Capilla-Lasheras, P., Dominoni, D.M., Babayan, S.A., O’Shaughnessy, P.J., Mladenova, M., Woodford, L., et al. (2017). Elevated Immune Gene Expression Is Associated with Poor Reproductive Success of Urban Blue Tits. Frontiers in Ecology and Evolution, 5, 64.

Cardilini, A.P.A., Weston, M.A., Nimmo, D.G., Dann, P. & Sherman, C.D.H. (2013). Surviving in sprawling suburbs: Suburban environments represent high quality breeding habitat for a widespread shorebird. Landscape and Urban Planning, 115, 72–80.

Chamberlain, D.E., Cannon, A.R., Toms, M.P. & Leech, D.I. (2009). Avian productivity in urban landscapes: a review and meta-analysis. Ibis, 151, 1–18.

Chao, A., Chiu, C. & Jost, L. (2014). Unifying Species Diversity, Phylogenetic Diversity, Functional Diversity, and Related Similarity and Differentiation Measures Through Hill Numbers. Annual Review of Ecology, Evolution, and Systematics, 45, 297–324.

Charter, M., Izhaki, I., Bouskila, A. & Leshem, Y. (2007). Breeding success of the Eurasian Kestrel (Falco tinnunculus) nesting on buildings in Israel. Journal of Raptor Research, 41, 139–143.

Cinar, O., Nakagawa, S. & Viechtbauer, W. (2022). Phylogenetic multilevel meta-analysis: A simulation study on the importance of modeling the phylogeny. Methods in Ecology and Evolution, 13, 383–395.

Cohen, J.E. & Xu, M. (2015). Random sampling of skewed distributions implies Taylor’s power law of fluctuation scaling. Proceedings of the National Academy of Science, 112, 7749–7754.

Conway, C.J., Garcia, V., Smith, M.D., Ellis, L.A. & Whitney, J.L. (2006). Comparative demography of Burrowing Owls in agricultural and urban landscapes in southeastern Washington. Journal of Field Ornithology, 77, 280–290.

Dhondt, A.A., Eyckerman, R., Moermans, R. & Hublé, J. (1984). Habitat and laying date of Great and Blue Tit Parus major and P. caeruleus. Ibis, 126, 388–397.

Diamond, S.E., Chick, L.D., Perez, A., Strickler, S.A. & Martin, R.A. (2018). Evolution of thermal tolerance and its fitness consequences: parallel and non-parallel responses to urban heat islands across three cities. Proceedings of the Royal Society B, 285, 20180036.

Diamond, S.E. & Martin, R.A. (2021). Evolution in Cities. Annual Review of Ecology, Evolution, and Systematics, 52, 519–540.

Dominoni, D., Quetting, M. & Partecke, J. (2013). Artificial light at night advances avian reproductive physiology. Proceedings of the Royal Society B, 280, 20123017.

Dunn, P.O. & Møller, A.P. (2014). Changes in breeding phenology and population size of birds. Journal of Animal Ecology, 83, 729–739.

Eden, S.F. (1985). The comparative breeding biology of magpies Pica pica in an urban and a rural habitat (Aves: Corvidae). Journal of Zoology, 205, 325–334.

ESA. 3CS Land Cover Product User Guide. (2020). ESA. Land Cover CCI Product User Guide. (2017).

Evans, B.A. & Gawlik, D.E. (2020). Urban food subsidies reduce natural food limitations and reproductive costs for a wetland bird. Scientific Reports, 10, 14021.

Fugère, V. & Hendry, A.P. (2018). Human influences on the strength of phenotypic selection. Proceedings of the National Academy of Science, 115, 10070–10075.

Fusco, G. (2001). How many processes are responsible for phenotypic evolution? Evolution & Development, 3, 279–286.

Fusco, N.A., Pehek, E. & Munshi-South, J. (2021). Urbanization reduces gene flow but not genetic diversity of stream salamander populations in the New York City metropolitan area. Evolutionary Applications, 14, 99–116.

Gahbauer, M.A., Bird, D.M., Clark, K.E., French, T., Brauning, D.W. & McMorris, F.A. (2015). Productivity, mortality, and management of urban peregrine falcons in northeastern North America. Journal of Wildlife Management, 79, 10–19.

Garant, D., Hadfield, J.D., Kruuk, L.E.B. & Sheldon, B.C. (2008). Stability of genetic variance and covariance for reproductive characters in the face of climate change in a wild bird population. Molecular Ecology, 17, 179–188.

Glądalski, M., Bańbura, M., Kaliński, A., Markowski, M., Skwarska, J., Wawrzyniak, J., et al. (2015). Inter-annual and inter-habitat variation in breeding performance of blue tits (Cyanistes caeruleus) in central Poland. Ornis Fennica, 92, 34–42.

Glądalski, M., Bańbura, M., Kaliński, A., Markowski, M., Skwarska, J., Wawrzyniak, J., et al. (2016a). Effects of extreme thermal conditions on plasticity in breeding phenology and double-broodedness of Great Tits and Blue Tits in central Poland in 2013 and 2014. International Journal of Biometeorology, 60, 1795–1800.

Glądalski, M., Bańbura, M., Kaliński, A., Markowski, M., Skwarska, J., Wawrzyniak, J., et al. (2016b). Effects of nest characteristics on reproductive performance in Blue Tits Cyanistes caeruleus and Great Tits Parus major. Avian Biology Research, 9, 37–43.

Glądalski, M., Bańbura, M., Kaliński, A., Markowski, M., Skwarska, J., Wawrzyniak, J., et al. (2017). Differences in the breeding success of blue tits Cyanistes caeruleus between a forest and an urban area: A long-term study. Acta Ornithologica, 52, 59–68.

Glądalski, M., Kaliński, A., Wawrzyniak, J., Bańbura, M., Markowski, M., Skwarska, J., et al. (2018). Physiological condition of nestling great tits Parus major in response to experimental reduction in nest micro- and macro-parasites. Conservation Physiology, 6, 1–9.

Gorton, A.J., Moeller, D.A. & Tiffin, P. (2018). Little plant, big city: A test of adaptation to urban environments in common ragweed (Ambrosia artemisiifolia). Proceedings of the Royal Society B, 285, 20180968.

Grafen, A. (1989). The phylogenetic regression. Philosophical Transactions of the Royal society of London, 326, 119–157.

Grimm, N.B., Faeth, S.H., Golubiewski, N.E., Redman, C.L., Wu, J., Bai, X., et al. (2008). Global change and the ecology of cities. Science, 319, 756–760.

Gryz, J. & Krauze-Gryz, D. (2018). Influence of habitat urbanisation on time of breeding and productivity of tawny Owl (Strix aluco). Polish Journal of Ecology, 66, 153–161.

Hajdasz, A.C., Otter, K.A., Baldwin, L.K. & Reudink, M.W. (2019). Caterpillar phenology predicts differences in timing of mountain chickadee breeding in urban and rural habitats. Urban Ecosystems, 22, 1113–1122.

Halfwerk, W., Blaas, M., Kramer, L., Hijner, N., Trillo, P.A., Bernal, X.E., et al. (2019). Adaptive changes in sexual signalling in response to urbanization. Nature Ecology and Evolution, 3, 374–380.

Hedges, L., Gurevitch, J. & Curtis, P. (1999). The Meta-Analysis of Response Ratios in Experimental Ecology. Ecology, 80, 1150–1156.

Hedges, L. V. (1981). Distribution theory for Glass’s estimator of effect size and related estimators. Journal of Educational Statistics, 6, 107–128.

Heisler, J.M. & Brazel, A.J. (2018). The urban physical environment: temperature and urban heat islands. In: Urban Ecosystem Ecology (eds. Aitkenhead-Petersn, J. & Volder, A.). American Society of Agronomy, Crop Science Society of America, Soil Science Society of America, Madison, pp. 29–56.

Hendry, A.P., Gotanda, K.M. & Svensson, E.I. (2017). Human influences on evolution, and the ecological and societal consequences. Philosophical Transactions of the Royal Society B, 372, 20160028.

Hijmans, R.J. (2020). raster: Geographic Data Analysis and Modeling.

Hill, M.O. (1973). Diversity and Evenness: A Unifying Notation and Its Consequences. Ecology, 54, 427–432.

Hinchliff, C.E., Smith, S.A., Allman, J.F., Burleigh, J.G., Chaudhary, R., Coghill, L.M., et al. (2015). Synthesis of phylogeny and taxonomy into a comprehensive tree of life. Proceedings of the National Academy of Science, 112, 12764–12769.

Hinsley, S.A., Hill, R.A., Bellamy, P.E., Harrison, N.M., Speakman, J.R., Wilson, A.K., et al. (2008). Effects of structural and functional habitat gaps on breeding woodland birds: Working harder for less. Landscape Ecology, 23, 615–626.

Ibáñez-Álamo, J.D. & Soler, M. (2010). Does urbanization affect selective pressures and life-history strategies in the common blackbird (Turdus merula L.)? Biological Journal of the Linnean Society, 101, 759–766.

Isaksson, C. & Andersson, S. (2007). Carotenoid diet and nestling provisioning in urban and rural great tits Parus major. Journal of Avian Biology, 38, 564–572.

Isaksson, C., Johansson, A. & Andersson, S. (2008). Egg yolk carotenoids in relation to habitat and reproductive investment in the great tit Parus major. Physiological and Biochemical Zoology, 81, 112–118.

Jarrett, C., Powell, L.L., McDevitt, H., Helm, B. & Welch, A.J. (2020). Bitter fruits of hard labour: diet metabarcoding and telemetry reveal that urban songbirds travel further for lower-quality food. Oecologia, 193, 377–388.

Johnson, M.T.J. & Munshi-South, J. (2017). Evolution of life in urban environments. Science, 358, eaam8327.

Kelleher, K.M. & O’Halloran, J. (2007). Influence of nesting habitat on breeding Song Thrushes Turdus philomelos. Bird Study, 54, 221–229.

Kettel, E.F., Gentle, L.K., Yarnell, R.W. & Quinn, J.L. (2019). Breeding performance of an apex predator, the peregrine falcon, across urban and rural landscapes. Urban Ecosystems, 22, 117–125.

Kopij, G. (2017). Changes in the number of nesting pairs and breeding success of the White Stork Ciconia ciconia in a large city and a neighbouring rural area in South-West Poland. Ornis Hungarica, 25, 109–115.

Lambert, M.R., Brans, K.I., Des Roches, S., Donihue, C.M. & Diamond, S.E. (2020). Adaptive Evolution in Cities: Progress and Misconceptions. Trends in Ecology and Evolution, 36, 239–257.

Lee, S., Lee, H., Jablonski, P.G., Choe, J.C. & Husby, M. (2017). Microbial abundance on the eggs of a passerine bird and related fitness consequences between urban and rural habitats. PLoS ONE, 12, e0185411.

Lin, W.L., Lin, S.M., Lin, J.W., Wang, Y. & Tseng, H.Y. (2015). Breeding performance of Crested Goshawk Accipiter trivirgatus in urban and rural environments of Taiwan. Bird Study, 62, 177–184.

De Lisle, S.P., Mäenpää, M.I. & Svensson, E.I. (2022). Phenotypic plasticity is aligned with phenological adaptation on both micro- and macroevolutionary timescales. Ecology Letters, 25, 790–801.

Liven-Schulman, I., Leshem, Y., Alon, D. & Yom-Tov, Y. (2004). Causes of population declines of the Lesser Kestrel Falco naumanni in Israel. Ibis, 146, 145–152.

Luna, Á., Palma, A., Sanz-Aguilar, A., Tella, J.L. & Carrete, M. (2020). Sex, personality and conspecific density influence natal dispersal with lifetime fitness consequences in urban and rural burrowing owls. PLoS ONE, 15, 1–17.

Luo, D., Wan, X., Liu, J. & Tong, T. (2016). Optimally estimating the sample mean from the sample size, median, mid-range, and/or mid-quartile range. Statistical Methods in Medical Research, 27, 1785–1805.

Mcgowan, K.J. (2001). Avian Ecology and Conservation in an Urbanizing World. In: Avian Ecology and Conservation in an Urbanizing World (eds. Marzluff, J.M., Bowman, R. & Donelly, R.). Kluwer Academic Press, New York, pp. 365–381.

Mennechez, G. & Clergeau, P. (2006). Effect of urbanisation on habitat generalists: starlings not so flexible? Acta Oecologica, 30, 182–191.

Merckx, T., Nielsen, M.E., Heliölä, J., Kuussaari, M., Pettersson, L.B., Pöyry, J., et al. (2021). Urbanization extends flight phenology and leads to local adaptation of seasonal plasticity in Lepidoptera. Proceedings of the National Academy of Sciences, 118, e2106006118.

Merckx, T., Souffreau, C., Kaiser, A., Baardsen, L.F., Backeljau, T., Bonte, D., et al. (2018). Body-size shifts in aquatic and terrestrial urban communities. Nature, 558, 113–116.

Michonneau, F., Brown, J.W. & Winter, D.J. (2016). rotl: an R package to interact with the Open Tree of Life data. Methods in Ecology and Evolution, 7, 1476–1481.

Middleton, A. (1979). Influence of Age and Habitat on Reproduction by the American Goldfinch. Ecology, 60, 418–432.

Millsap, B., Breen, T., McConnell, E., Steffer, T., Phillips, L., Douglass, N., et al. (2004). Comparative Fecundity and Survival of Bald Eagles Fledged from Suburban and Rural Natal Areas in Florida. Journal of Wildlife Management, 68, 1018–1031.

Minias, P. (2016). Reproduction and survival in the city: Which fitness components drive urban colonization in a reed-nesting waterbird? Current Zoology, 62, 79–87.

Morrissey, C.A., Stanton, D.W.G., Tyler, C.R., Pereira, M.G., Newton, J., Durance, I., et al. (2014). Developmental impairment in eurasian dipper nestlings exposed to urban stream pollutants. Environmental Toxicology and Chemistry, 33, 1315–1323.

Muñoz, M.M., Stimola, M.A., Algar, A.C., Conover, A., Rodriguez, A.J., Landestoy, M.A., et al. (2014). Evolutionary stasis and lability in thermal physiology in a group of tropical lizards. Proceedings of the Royal Society B, 281, 20132433.

Nakagawa, S., Lagisz, M., Jennions, M.D., Koricheva, J., Noble, D.W.A., Parker, T.H., et al. (2022). Methods for testing publication bias in ecological and evolutionary meta-analyses. Methods in Ecology and Evolution, 13, 4–21.

Nakagawa, S., Lagisz, M., O’Dea, R.E., Rutkowska, J., Yang, Y., Noble, D.W.A., et al. (2021). The orchard plot: Cultivating a forest plot for use in ecology, evolution, and beyond. Research Synthesis Methods, 12, 4–12.

Nakagawa, S., Poulin, R., Mengersen, K., Reinhold, K., Engqvist, L., Lagisz, M., et al. (2015). Meta-analysis of variation: Ecological and evolutionary applications and beyond. Methods in Ecology and Evolution, 6, 143–152.

Nakagawa, S. & Santos, E.S.A. (2012). Methodological issues and advances in biological meta-analysis. Evolutionary Ecology, 26, 1253–1274.

Nakagawa, S. & Schielzeth, H. (2013). A general and simple method for obtaining R2 from generalized linear mixed-effects models. Methods in Ecology and Evolution, 4, 133–142.

Newhouse, M.J., Marra, P.P. & Johnson, L.S. (2008). Reproductive Success of House Wrens in Suburban and Rural Landscapes. The Wilson Journal of Ornithology, 120, 99–104.

OpenTreeOfLife, Redelings, R., Reyes, L.L.S., Cranston, K.A., Allman, J., Holder, M.T., et al. (2019). Open Tree of Life Synthetic Tree.

Partecke, J., Hegyi, G., Fitze, P.S., Gasparini, J. & Schwabl, H. (2020). Maternal effects and urbanization: Variation of yolk androgens and immunoglobulin in city and forest blackbirds. Ecology and Evolution, 10, 2213–2224.

Pavlicev, M., Cheverud, J.M. & Wagner, G.P. (2011). Evolution of adaptive phenotypic variation patterns by direct selection for evolvability. Proceedings of the Royal Society B, 278, 1903–1912.

Pebesma, E. (2018). Simple Features for R: Standardized Support for Spatial Vector Data. The R Journal, 10, 439–446.

Perlut, N.G., Bonter, D.N., Ellis, J.C. & Friar, M.S. (2016). Roof-Top Nesting in a Declining Population of Herring Gulls (Larus argentatus) in Portland, Maine, USA. Waterbirds, 39, 68–73.

Pickett, S.T.A., Cadenasso, M.L., Rosi-Marshall, E.J., Belt, K.T., Groffman, P.M., Grove, J.M., et al. (2017). Dynamic heterogeneity: a framework to promote ecological integration and hypothesis generation in urban systems. Urban Ecosystems, 20, 1–14.

Pollock, C.J., Capilla-Lasheras, P., McGill, R.A.R., Helm, B. & Dominoni, D.M. (2017). Integrated behavioural and stable isotope data reveal altered diet linked to low breeding success in urban-dwelling blue tits (Cyanistes caeruleus). Scientific Reports, 7, 5014.

Preiszner, B., Papp, S., Pipoly, I., Seress, G., Vincze, E., Liker, A., et al. (2017). Problem-solving performance and reproductive success of great tits in urban and forest habitats. Animal Cognition, 20, 53–63.

R Core Team. (2022). R: A language and environment for statistical computing. R Foundation for Statistical Computing, Vienna, Austria.

Rees, J.A. & Cranston, K. (2017). Automated assembly of a reference taxonomy for phylogenetic data synthesis. Biodiversity Data Journal, 5, e12581.

Richardson, J.L., Michaelides, S., Combs, M., Djan, M., Bisch, L., Barrett, K., et al. (2021). Dispersal ability predicts spatial genetic structure in native mammals persisting across an urbanization gradient. Evolutionary Applications, 14, 163–177.

Rollinson, D.J. & Jones, D.N. (2003). Variation in breeding parameters of the Australian magpie Gymnorhina tibicen in suburban and rural environments. Urban Ecosystems, 6, 257–269.

Rosenfield, R.N., Hardin, M.G., Taylor, J., Sobolik, L.E. & Frater, P.N. (2019). Nesting density and dispersal movements between urban and rural habitats of cooper’s Hawks (Accipiter cooperii) in Wisconsin: Are these source or sink habitats? American Midland Naturalist, 182, 36–51.

Rowe, L., Ludwig, D. & Schluter, D. (1994). Time, condition, and the seasonal decline of avian clutch size. American Naturalist, 143, 698–772.

Salmón, P., Jacobs, A., Ahrén, D., Biard, C., Dingemanse, N.J., Dominoni, D.M., et al. (2021). Continent-wide genomic signatures of adaptation to urbanisation in a songbird across Europe. Nature Communications, 12, 2983.

Santangelo, J.S., Ness, R.W., Cohan, B., Fitzpatrick, C.R., Innes, S.G., Koch, S., et al. (2022). Global urban environmental change drives adaptation in white clover. Science, 375, 1275–1281.

de Satgé, J., Strubbe, D., Elst, J., De Laet, J., Adriaensen, F. & Matthysen, E. (2019). Urbanisation lowers great tit Parus major breeding success at multiple spatial scales. Journal of Avian Biology, 50, e02108.

Schmidt, V.K.-H. & Steinbach, J. (1983). Niedriger Bruterfolg der Kohlmeise (Parus major) in städtischen Parks und Friedhöfen. Journal für Ornithologie, 124, 81–83.

Schoech, S.J. & Bowman, R. (2001). Variation in the timing of breeding between suburban and wildland Florida Scrub-Jays: Do physiologic measures reflect different environments? In: Avian Ecology and Conservation in an Urbanizing World (eds. Marzluff, J.M., Bowman, R. & Donelly, R.). Kluwer Academic Press, New York, pp. 289–306.

Schoech, S.J., Bowman, R., Bridge, E.S. & Boughton, R.K. (2007). Baseline and acute levels of corticosterone in Florida Scrub-Jays (Aphelocoma coerulescens): Effects of food supplementation, suburban habitat, and year. General and Comparative Endocrinology, 154, 150–160.

Schoech, S.J., Bowman, R. & Reynolds, S.J. (2004). Food supplementation and possible mechanisms underlying early breeding in the Florida Scrub-Jay (Aphelocoma coerulescens). Hormones and Behavior, 46, 565–573.

Schoech, S.J., Bridge, E.S., Boughton, R.K., Reynolds, S.J., Atwell, J.W. & Bowman, R. (2008). Food supplementation: A tool to increase reproductive output? A case study in the threatened Florida Scrub-Jay. Biological Conservation, 141, 162–173.

Senior, A.M., Gosby, A.K., Lu, J., Simpson, S.J. & Raubenheimer, D. (2016a). Meta-analysis of variance: An illustration comparing the effects of two dietary interventions on variability in weight. Evolution, Medicine and Public Health, 2016, 244–255.

Senior, A.M., Grueber, C.E., Kamiya, T., Lagisz, M., O’Dwyer, K., Santos, E.S.A., et al. (2016b). Heterogeneity in ecological and evolutionary meta-analyses: its magnitude and implications. Ecology, 97, 3293–3299.

Senior, A.M., Viechtbauer, W. & Nakagawa, S. (2020). Revisiting and expanding the meta-analysis of variation: The log coefficient of variation ratio. Research Synthesis Methods, 11, 553–567.

Sepp, T., McGraw, K.J., Kaasik, A. & Giraudeau, M. (2018). A review of urban impacts on avian life-history evolution: Does city living lead to slower pace of life? Global Change Biology, 24, 1452–1469.

Seress, G., Bókony, V., Pipoly, I., Szép, T., Nagy, K. & Liker, A. (2012). Urbanization, nestling growth and reproductive success in a moderately declining house sparrow population. Journal of Avian Biology, 43, 403–414.

Seress, G., Hammer, T., Bókony, V., Vincze, E., Preiszner, B., Pipoly, I., et al. (2018). Impact of urbanization on abundance and phenology of caterpillars and consequences for breeding in an insectivorous bird. Ecological Applications, 28, 1143–1156.

Seress, G., Sándor, K., Evans, K.L. & Liker, A. (2020). Food availability limits avian reproduction in the city: An experimental study on great tits Parus major. Journal of Animal Ecology, 89, 1570–1580.

Sharma, R.C., Bhatt, D. & Sharma, R.K. (2004). Breeding success of the tropical Spotted Munia Lonchura punctulata in urbanized and forest habitats. Ornithological Science, 3, 113–117.

Sheldon, B.C., Kruuk, L.E.B. & Merilä, J. (2003). Natural Selection and Inheritance of Breeding Time and Clutch Size in the Collared Flycatcher. Evolution, 57, 406–420.

Shi, J., Luo, D., Weng, H., Zeng, X., Lin, L., Chu, H., et al. (2020). Optimally estimating the sample standard deviation from the five-number summary. Research Synthesis Methods, 11, 641–654.

Shochat, E., Warren, P.S., Faeth, S.H., McIntyre, N.E. & Hope, D. (2006). From patterns to emerging processes in mechanistic urban ecology. Trends in Ecology and Evolution, 21, 186–191.

Shustack, D.P. & Rodewald, A.D. (2011). Nest predation reduces benefits to early clutch initiation in northern cardinals Cardinalis cardinalis. Journal of Avian Biology, 42, 204–209.

Solonen, T. (2001). Breeding of the Great Tit and Blue Tit in urban and rural habitats in Southern Finland. Ornis Fennica, 78, 49–60.

Solonen, T. (2014). Timing of breeding in rural and urban Tawny Owls Strix aluco in southern Finland: Effects of vole abundance and winter weather. Journal of Ornithology, 155, 27–36.

Solonen, T. & Ursin, K.A. (2008). Breeding of Tawny Owls Strix aluco in rural and urban habitats in southern Finland. Bird Study, 55, 216–221.

Stout, W.E., Anderson, R.K. & Papp, J.M. (1998). Urban, suburban and rural red-tailed hawk nesting habitat and populations in southeast Wisconsin. Journal of Raptor Research, 32, 221–228.

Stracey, C.M. & Robinson, S.K. (2012). Are urban habitats ecological traps for a native songbird? Season-long productivity, apparent survival, and site fidelity in urban and rural habitats. Journal of Avian Biology, 43, 50–60.

Strubbe, D., Salleh Hudin, N., Teyssier, A., Vantieghem, P., Aerts, J. & Lens, L. (2020). Phenotypic signatures of urbanization are scale-dependent: A multi-trait study on a classic urban exploiter. Landscape and Urban Planning, 197, 103767.

Sumasgutner, P., Nemeth, E., Tebb, G., Krenn, H.W. & Gamauf, A. (2014). Hard times in the city-attractive nest sites but insufficient food supply lead to low reproduction rates in a bird of prey. Frontiers in Zoology, 34, 17–31.

Thompson, M.J., Capilla-Lasheras, P., Dominoni, D.M., Réale, D. & Charmantier, A. (2022). Phenotypic variation in urban environments: mechanisms and implications. Trends in Ecology & Evolution, 37, 171–182.

Thornton, M., Todd, I. & Roos, S. (2017). Breeding Success and Productivity of Urban and Rural Eurasian Sparrowhawks Accipiter nisus in Scotland. Ecoscience, 24, 115–116.

Trikalinos, T. & Ioannidis, J.P.A. (2005). Assessing the evolution of effect sizes over time. In: Publication Bias in Meta-Analysis (eds. Rothstein, H., Sutton, A. & Borenstein, M.). John Wiley, Chichester, pp. 241–259.

Uchida, K., Blakey, R. V., Burger, J.R., Cooper, D.S., Niesner, C.A. & Blumstein, D.T. (2021). Urban Biodiversity and the Importance of Scale. Trends in Ecology and Evolution, 36, 123–131.

Via, S. & Lande, R. (1985). Genotype-Environment Interaction and the Evolution of Phenotypic Plasticity. Evolution, 39, 505–522.

Viechtbauer, W. (2010). Conducting meta-analyses in R with the metafor. Journal of Statistical Software, 36, 1–48.

Wawyrzyniak, J., Kaliński, A., Glądalski, M., Bańbura, M., Markowski, M., Skwarska, J., et al. (2015). Long-Term Variation in Laying Date and Clutch Size of the Great Tit Parus major in Central Poland: A Comparison between Urban Parkland and Deciduous Forest. Ardeola, 62, 311–322.

Welch-Acosta, B.C., Skipper, B.R. & Boal, C.W. (2019). Comparative breeding ecology of Mississippi Kites in urban and exurban areas of West Texas. Journal of Field Ornithology, 90, 248–257.

Westgate, M.J. (2019). revtools: An R package to support article screening for evidence synthesis. Research Synthesis Methods, 10, 606–614.

Yeh, P.J. & Price, T.D. (2004). Adaptive phenotypic plasticity and the successful colonization of a novel environment. American Naturalist, 164, 531–542.

