## Supplementary materials for "A global meta-analysis reveals higher variation in breeding phenology in urban birds than in their non-urban neighbours"

**This file contains Supplementary figures, Supplementary tables, and Supplementary text.**

### Supplementary figures & tables

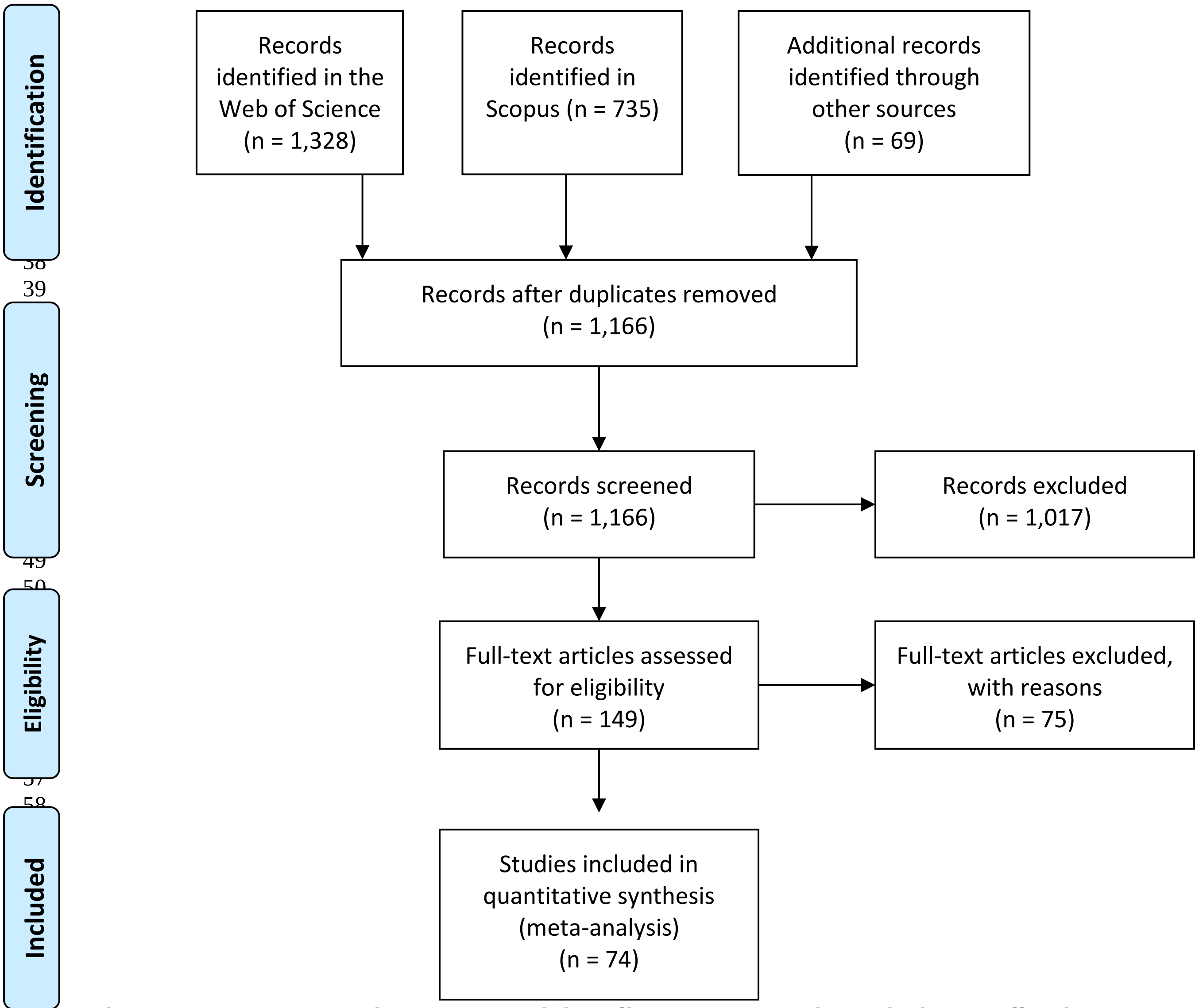


**Figure S1. PRISMA chart summarising literature search and data collection**. Potential papers with relevant information in Chamberlain *et al*. 2009 were identified from Appendix S1 (sections 'lay date', 'clutch size', 'fledglings per successful breeding attempt' and 'fledglings per attempt'; n = 37). In Sepp *et al.* 2018, we screened papers shown in Supplementary Materials Section 4, Table S5 and Table S14 (n = 32). Additionally, 1,328 and 735 published studies were identified through several searches in the Web of Science Core Collection and Scopus (see main text methods for details). Diagram template downloaded from: http://prisma-statement.org/prismastatement/flowdiagram.aspx

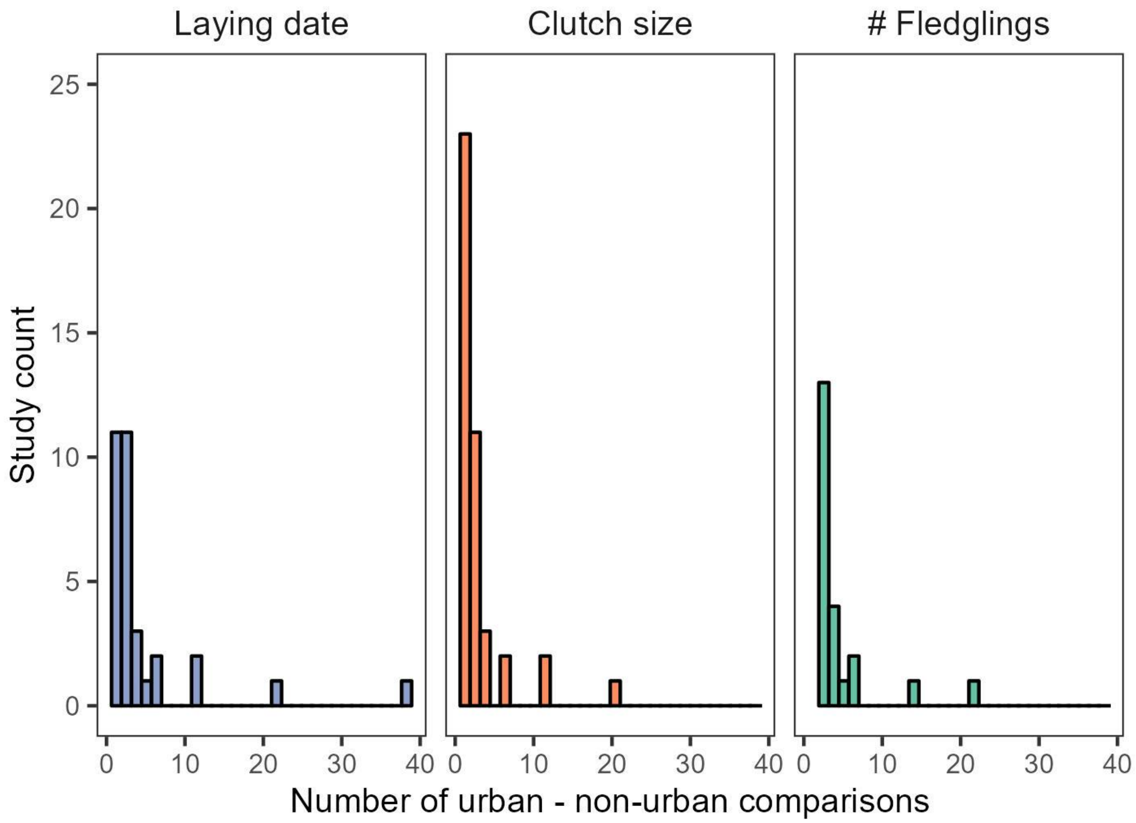


**Figure S2. Sample size distribution per trait**. Histograms represent the number of published studies (y axis) that reported a given number of urban – non-urban comparisons (x axis; e.g., for several breeding seasons) for each trait.

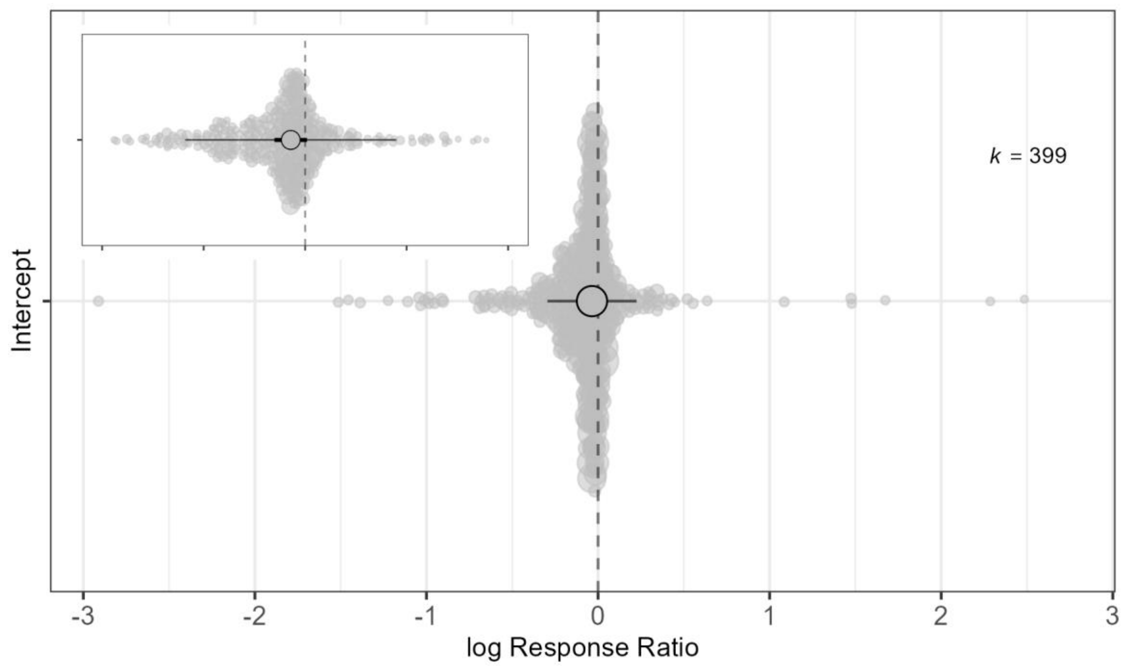


**Figure S3. Meta-analysis of differences in the log response ratio (lnRR) between urban and non-urban populations.** Estimates for laying dates, clutch size and number of fledglings per clutch are combined in this analysis (k = 399 observations). The snippet amplifies the figure between x = -0.5 and x = 0.5 to help visualise the overall effect size and its 95% confidence and prediction intervals. Positive values in the x axis represent higher mean values in urban than in non-urban populations, whereas negative values illustrate lower mean values in urban compared to paired non-urban populations. A vertical dashed line is drawn at an x value of zero. The large grey point, thick and thin intervals provide the model estimate, 95% confidence and prediction intervals respectively for the overall value of the log response ratio. Transparent small grey points show raw data (their size is scaled to illustrate the sample size from which they were estimated – e.g., the larger the point, the bigger the sample size) and ‘k’ provides the number of observations.

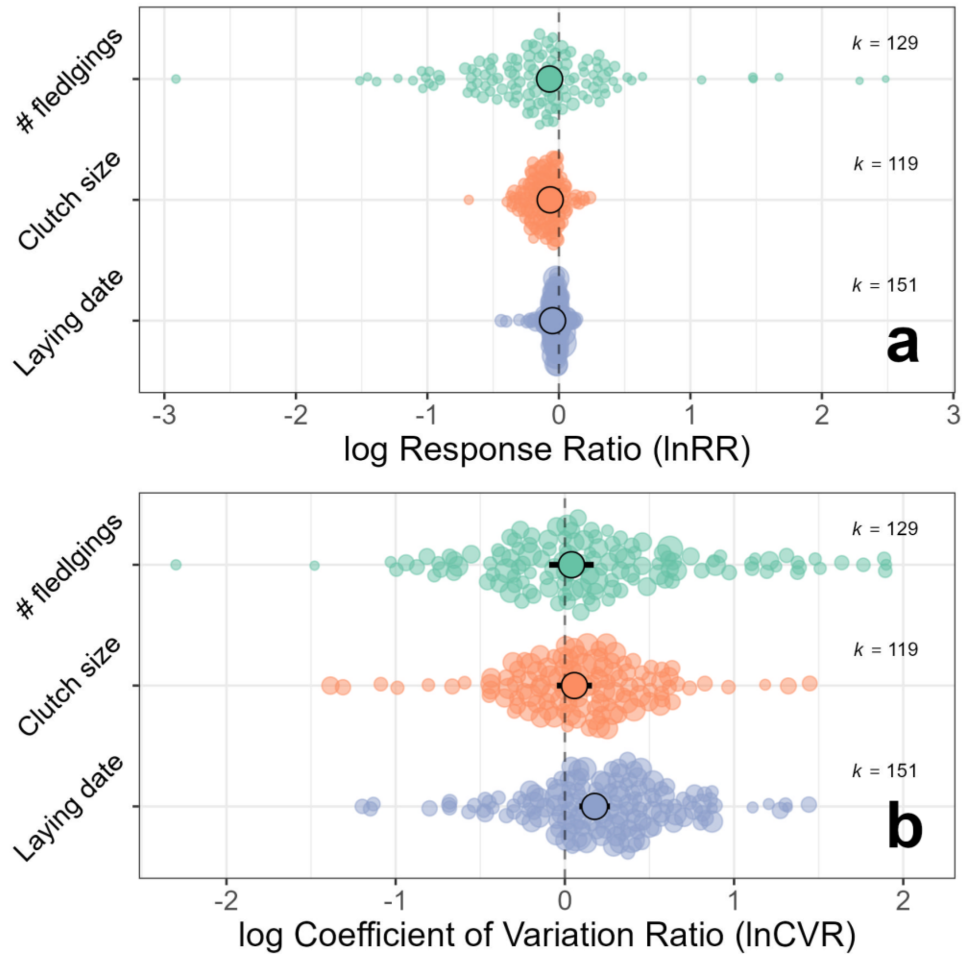


**Figure S4. Urban populations have earlier phenology, lower reproductive output and more variable life-history traits than non-urban populations**. (**a**) Urban populations laid earlier and had smaller clutches, producing smaller broods, than their paired non-urban populations (illustrated by negative lnRR estimates; Model 2). (**b**) Our meta-analysis revealed that variation in life-history traits was higher in urban populations compared to non-urban counterparts, with a marked difference between habitats in laying dates (illustrated by positive estimates of lnCVR, Model 4). Model estimates are shown along with their 95% confidence intervals per trait as calculated by Model 2 and 4 (see Table S3 & Table S5 for full model outputs). Transparent points illustrate raw data (the size of the point is scaled to illustrate the sample size from which they were estimated) and 'k' provides the number of urban – non-urban comparisons.

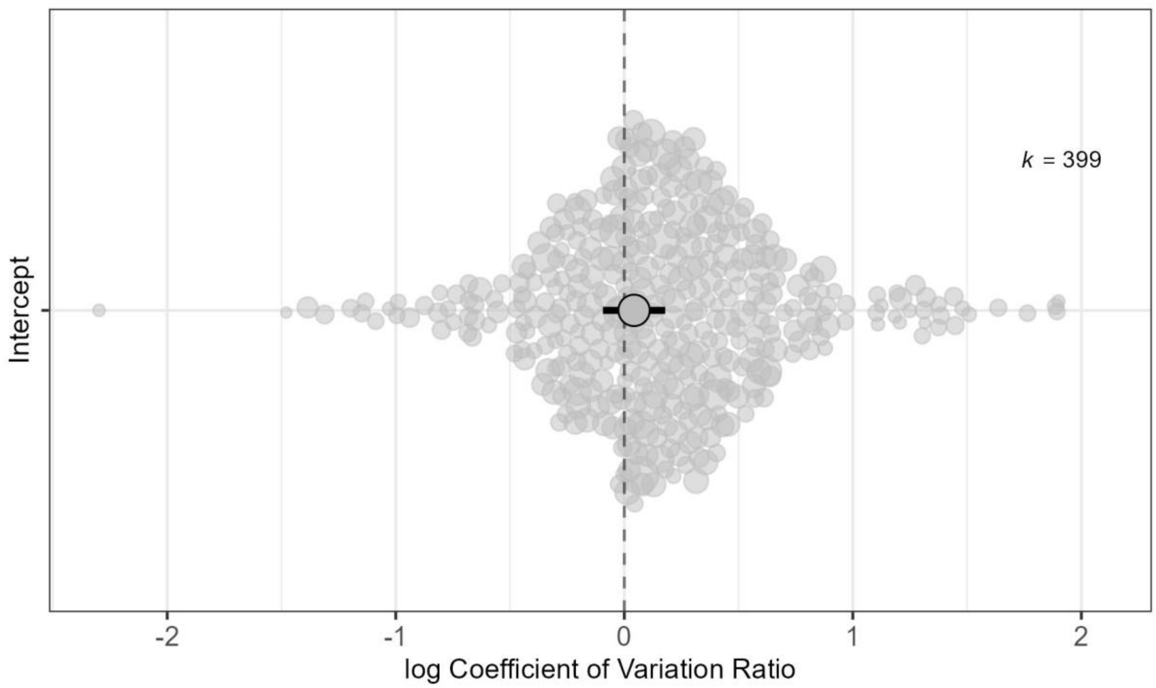


**Figure S5. Meta-analysis of differences in the log coefficient of variation ratio (lnCVR) between urban and non-urban populations.** Estimates for laying dates, clutch size and number of fledglings per clutch are combined in this initial analysis (k = 399 observations). Positive values in the x axis represent higher coefficient of variation (i.e., SD/mean$^2$) in urban populations than in non-urban populations, whereas negative values illustrate higher coefficient of variation in non-urban populations compared to paired urban populations. Vertical dashed line drawn at an x value of zero. The large grey point, thick and thin intervals provide the model estimate, 95% confidence and prediction intervals, respectively, for the overall value of the log of the coefficient of variation ratio. Transparent small grey points show raw data (their size is scaled to illustrate the sample size from which they were estimated – e.g., the larger the point, the bigger the sample size) and 'k' provides the number of observations.

#### Supplementary Tables

**Table S1.** Total (relative) heterogeneity (i.e., $I^2_{total}$, percentage of variation remaining after accounting for sampling variance), percentage of residual variation (i.e., $I^2_{observation\ ID}$) and the percentage of variation explained by each random term included in our multilevel meta-analyses of lnRR (Model 1) and lnCVR (Model 3). For more details on models see Table 1.

| Model ID | Response variable | $I^2_{total}$ | $I^2_{study\ ID}$ | $I^2_{Population\ ID}$ | $I^2_{phylogeny}$ | $I^2_{species\ ID}$ | $I^2_{observation\ ID}$ |
|---|---|---|---|---|---|---|---|
| Model 1 | lnRR | 97.8 | 8.4 | 12.3 | 1.7 | 15.5 | 59.9 |
| Model 3 | lnCVR | 74.3 | 0.0 | 11.6 | 5.8 | 3.3 | 53.6 |

**Table S2**. Statistical support for models testing differences in mean values of life-history traits between urban and non-urban populations (i.e., lnRR) employing different variance-covariance matrix structures to model correlations among traits (see Methods). 'UN': Unstructured variance-covariance matrix; 'HCS': Heteroscedastic compound symmetric variance-covariance matrix; 'DIAG': diagonal variance-covariance matrix; 'CS': Compound symmetric variance-covariance matrix; 'NONE': Single variance across traits and no correlations; g: number of model parameters. See section 'Methods' for model details. We refer to the top model in this table as 'Model 2'. Full model estimates in Table S3.

| Variance-covariance structure | g | Log-likelihood | AIC | ΔAIC |
|---|---|---|---|---|
| UN (Model 2) | 15 | 279.80 | -529.59 | 0 |
| HCS | 13 | 262.79 | -499.58 | 30.01 |
| DIAG | 12 | 261.78 | -499.56 | 30.03 |
| CS | 11 | 252.88 | -483.75 | 45.84 |
| NONE | 10 | 207.27 | -394.48 | 135.11 |

**Table S3.** Meta-analytic model (Model 2) coefficients explaining variation in lnRR (i.e., differences in mean life-history traits between urban and non-urban populations) allowing for different random effect variance-covariance for 'study ID' and 'observation ID' variance across traits (i.e., each trait has a [correlated] 'study ID' variance component and 'observation ID' variance). This model implemented the variance-covariance matrix structure of the top model in Table S2. CI = confidence interval; k = number of observations.

**Fixed Effects**

| | estimate | 95% CI | |
|---|---|---|---|
| Laying date | -0.048 | -0.084 | -0.012 |
| Clutch size | -0.066 | -0.107 | -0.025 |
| Number of fledglings | -0.070 | -0.171 | 0.032 |

**Random effect & residual variances**

| | Trait | estimate | k |
|---|---|---|---|
| Population ID | Laying date<br>Clutch size<br>Number of fledglings | 0.001 | 57 |
| Phylogeny | Laying date<br>Clutch size<br>Number of fledglings | 0 | 35 |
| Permanent species effect | Laying date<br>Clutch size<br>Number of fledglings | 0.003 | 35 |
| Study ID | Laying date | 0.003 | 151 |
| | Clutch size | 0.010 | 119 |
| | Number of fledglings | 0.098 | 129 |
| Study ID (correlations) | Laying date - Clutch size | -0.959 | |
| | Laying date - Number of fledglings | -0.843 | |
| | Clutch size - Number of fledglings | 0.859 | |
| Observation ID | Laying date | 0.001 | 151 |
| | Clutch size | 0.002 | 119 |
| | Number of fledglings | 0.036 | 129 |

**Table S4.** Statistical support for models testing differences in variation of life-history traits between urban and non-urban populations (i.e., lnCVR) employing different variance-covariance matrix structures to model correlations among traits (see Methods). 'DIAG': diagonal variance-covariance matrix; 'HCS': Heteroscedastic compound symmetric variance-covariance matrix; 'UN': Unstructured variance-covariance matrix; 'CS': Compound symmetric variance-covariance matrix; 'NONE': Single variance across traits and no correlations; k: number of model parameters. See section 'Methods' for model details. We refer to the top model in this table as 'Model 4'. Full model estimates in Table S4.

| Variance-covariance structure | g | Log-likelihood | AIC | ΔAIC |
|---|---|---|---|---|
| DIAG (Model 4) | 12 | -230.31 | 485.63 | 0 |
| HCS | 13 | -227.86 | 485.71 | 1.08 |
| UN | 15 | -230.17 | 486.34 | 1.71 |
| CS | 11 | -232.92 | 487.84 | 3.21 |
| NONE | 10 | -250.04 | 520.07 | 35.45 |

**Table S5.** Meta-analytic model (Model 4) coefficients explaining variation in lnCVR (i.e., differences in coefficient of variation in life-history traits between urban and non-urban bird populations) allowing for different random effect variance for ‘study ID’ and residual variance across traits (i.e., each trait has an independent ‘study ID’ variance and residual variance). This model implemented the variance-covariance matrix structure of the top model in Table S4. CI = confidence interval; k = number of observations.

**Fixed effects**

| | estimate | 95% CI | |
|---|---|---|---|
| Laying date | 0.176 | 0.084 | 0.268 |
| Clutch size | 0.055 | -0.051 | 0.160 |
| Number of fledglings | 0.037 | -0.096 | 0.171 |

**Random effect & residual variances**

| | Trait | estimate | k |
|---|---|---|---|
| Population ID | Laying date<br>Clutch size<br>Number of fledglings | 0 | 57 |
| Phylogeny | Laying date<br>Clutch size<br>Number of fledglings | 0 | 35 |
| Permanent species effect | Laying date<br>Clutch size<br>Number of fledglings | 0.006 | 35 |
| Study ID | Laying date | 0.010 | 151 |
| | Clutch size | 0.052 | 119 |
| | Number of fledglings | 0.124 | 129 |
| Observation ID | Laying date | 0.052 | 151 |
| | Clutch size | 0.030 | 119 |
| | Number of fledglings | 0.058 | 129 |

### Supplementary text

#### A – Assessment of systematic literature search

First, we used 31 papers found in the reference lists of Chamberlain *et al.* 2009 and Sepp *et al*. 2018, and that were included in our original meta-analytic database (before removing effect sizes with missing information) to assess the comprehensiveness of our literature search. Out of these 31 studies, 24 (77.4%) were found by our searches on the Web of Science Core Collection, while 22 (71.0%) were found by our search on Scopus (see details in the main text). All 22 papers found on Scopus were also found on the Web of Science Core Collection. 22 out of 24 papers found on the Web of Science Core Collection were also retrieved from Scopus, representing a 91.67% overlap between the two search engines. Second, we investigated the comprehensiveness of our literature search by comparing the search results from the Web of Science Core Collection and Scopus. Our most inclusive search string ('(1)' – see main text) returned 735 papers on Scopus. Out of these, 509 papers (69.3% overlap) had already been found through our searches on Web of Science Core Collection.

#### B – Data extraction validation

All effect sizes were extracted by one author (PC-L). To validate data extraction, another author (MJT) blindly checked 10 studies, 15% of the studies included the meta-analysis, comprising 55 effect sizes (17.8% of the 399 effect sizes included in the final dataset). The same number of effect sizes per study were extracted by both authors in 8 out of 10 cases. In one study, the data validator extracted no effect sizes but correctly identified that three effect sizes (those present in the final dataset) could be extracted either from figures presented in the paper or by contacting the authors of the study. In another study, the data validator extracted two out of three effect sizes present in the final dataset but correctly identified that an additional effect size could be extracted either from figures presented in the paper or by contacting the authors of the study. In the 55 effect sizes that both authors independently extracted, we found a high correlation in extracted mean values (Pearson's correlation coefficient between original data extraction and re-extraction: r [95% CI] = 0.997 [0.996, 0.998]), extracted standard deviations (r [95% CI] = 0.880 [0.828, 0.917]) and extracted sample sizes (r [95% CI] = 0.998 [0.998, 0.999]). The slightly lower r value for standard deviation was entirely due to estimates from one study (Dhondt *et al.* 1984), in which the authors extracted data from different urban populations. Excluding this study, the correlation between extracted standard deviations was virtually one (r [95%CI] = 0.999 [0.999,0.999]). Our dataset contains a column providing details on data extraction for every single effect size (see dataset in GitHub repository: https://github.com/PabloCapilla/meta-analysis_variation_urban) and, therefore, reproducing our data extraction process should yield repeatable results, as shown above.

#### C – Assessment of publication bias

We found little evidence for the existence of small-study or decline effects for lnRR (slope estimate for the square-root of the inverse of the effective sample size [95% CI] = 0.041 [-0.096, 0.178], $R^2_{margnal}$ = 0.25%; slope estimate for year of publication [95% CI] = 0.001 [-0.003, 0.003], $R^2_{margnal}$ = 0.02%) or lnCVR (slope estimate for the square-root of the inverse of the effective sample size [95% CI] = 0.241 [-0.174, 0.655], $R^2_{margnal}$ = 1.21%; slope estimate for year of publication [95% CI] = -0.001 [-0.008, 0.006], $R^2_{margnal}$ = 0.05%). These findings suggest that our results do not seemingly suffer from publications bias (Nakagawa *et al.* 2022).

#### D – Search string including a list of all avian genera

The string below is presented in the format required for a search on the Web of Science. The string was adjusted to carry out the search on Scopus.

TS=("urban*" AND ("bird*" OR "aves" OR "avian" OR "ornithol*" OR "passerine*" OR "passeriform*" OR "songbird*" OR "Accipiter" OR "Aegypius" OR "Aquila" OR "Aviceda" OR "Busarellus" OR "Butastur" OR "Buteo" OR "Buteogallus" OR "Chondrohierax" OR "Circaetus" OR "Circus" OR "Clanga" OR "Elanoides" OR "Elanus" OR "Gampsonyx" OR "Geranoaetus" OR "Geranospiza" OR "Gypaetus" OR "Gypohierax" OR "Gyps" OR "Haliaeetus" OR "Haliastur" OR "Harpagus" OR "Harpia" OR "Hieraaetus" OR "Ictinaetus" OR "Ictinia" OR "Kaupifalco" OR "Leptodon" OR "Leucopternis" OR "Lophotriorchis" OR "Milvus" OR "Morphnarchus" OR "Morphnus" OR "Necrosyrtes" OR "Neophron" OR "Nisaetus" OR "Parabuteo" OR "Pernis" OR "Pithecophaga" OR "Polemaetus" OR "Pseudastur" OR "Rostrhamus" OR "Rupornis" OR "Sarcogyps" OR "Spilornis" OR "Spizaetus" OR "Stephanoaetus" OR "Terathopius" OR "Torgos" OR "Trigonoceps" OR "Cathartes" OR "Coragyps" OR "Gymnogyps" OR "Sarcoramphus" OR "Vultur" OR "Pandion" OR "Sagittarius" OR "Aix" OR "Alopochen" OR "Amazonetta" OR "Anas" OR "Anser" OR "Asarcornis" OR "Aythya" OR "Branta" OR "Bucephala" OR "Cairina" OR "Callonetta" OR "Cereopsis" OR "Chen" OR "Chenonetta" OR "Chloephaga" OR "Clangula" OR "Coscoroba" OR "Cyanochen" OR "Cygnus" OR "Dendrocygna" OR "Heteronetta" OR "Histrionicus" OR "Hymenolaimus" OR "Lophodytes" OR "Lophonetta" OR "Malacorhynchus" OR "Mareca" OR "Marmaronetta" OR "Melanitta" OR "Merganetta" OR "Mergellus" OR "Mergus" OR "Neochen" OR "Netta" OR "Nettapus" OR "Nomonyx" OR "Oxyura" OR "Plectropterus" OR "Polysticta" OR "Pteronetta" OR "Sarkidiornis" OR "Sibirionetta" OR "Somateria" OR "Spatula" OR "Tachyeres" OR "Tadorna" OR "Thalassornis" OR "Anhima" OR "Chauna" OR "Anseranas" OR "Aegotheles" OR "Aerodramus" OR "Aeronautes" OR "Apus" OR "Chaetura" OR "Collocalia" OR "Cypseloides" OR "Cypsiurus" OR "Hirundapus" OR "Panyptila" OR "Streptoprocne" OR "Tachornis" OR "Tachymarptis" OR "Hemiprocne" OR "Abeillia" OR "Adelomyia" OR "Aglaeactis" OR "Aglaiocercus" OR "Amazilia" OR "Androdon" OR "Anopetia" OR "Anthocephala" OR "Anthracothorax" OR "Archilochus" OR "Atthis"

OR "Augastes" OR "Avocettula" OR "Basilinna" OR "Boissonneaua" OR "Calliphlox" OR "Calypte" OR "Campylopterus" OR "Chaetocercus" OR "Chalcostigma" OR "Chalybura" OR "Chlorestes" OR "Chlorostilbon" OR "Chrysolampis" OR "Chrysuronia" OR "Clytolaema" OR "Coeligena" OR "Colibri" OR "Cyanophaia" OR "Cynanthus" OR "Discosura" OR "Doricha" OR "Doryfera" OR "Elvira" OR "Ensifera" OR "Eriocnemis" OR "Eugenes" OR "Eulampis" OR "Eulidia" OR "Eupetomena" OR "Eupherusa" OR "Eutoxeres" OR "Florisuga" OR "Glaucis" OR "Goethalsia" OR "Goldmania" OR "Haplophaedia" OR "Heliactin" OR "Heliangelus" OR "Heliodoxa" OR "Heliomaster" OR "Heliothryx" OR "Hylocharis" OR "Juliamyia" OR "Klais" OR "Lafresnaya" OR "Lampornis" OR "Lamprolaima" OR "Lepidopyga" OR "Lesbia" OR "Leucippus" OR "Leucochloris" OR "Lophornis" OR "Mellisuga" OR "Metallura" OR "Microchera" OR "Myrmia" OR "Myrtis" OR "Ocreatus" OR "Opisthoprora" OR "Oreonympha" OR "Oreotrochilus" OR "Orthorhyncus" OR "Oxypogon" OR "Panterpe" OR "Patagona" OR "Phaeochroa" OR "Phaethornis" OR "Phlogophilus" OR "Polyonymus" OR "Polytmus" OR "Pterophanes" OR "Ramphodon" OR "Ramphomicron" OR "Rhodopis" OR "Sappho" OR "Schistes" OR "Selasphorus" OR "Sephanoides" OR "Stephanoxis" OR "Sternoclyta" OR "Taphrospilus" OR "Thalurania" OR "Thaumastura" OR "Threnetes" OR "Topaza" OR "Trochilus" OR "Urochroa" OR "Urosticte" OR "Apteryx" OR "Anthracoceros" OR "Buceros" OR "Bycanistes" OR "Lophoceros" OR "Penelopides" OR "Rhabdotorrhinus" OR "Rhyticeros" OR "Tockus" OR "Bucorvus" OR "Phoeniculus" OR "Rhinopomastus" OR "Upupa" OR "Antrostomus" OR "Caprimulgus" OR "Chordeiles" OR "Eleothreptus" OR "Eurostopodus" OR "Hydropsalis" OR "Lurocalis" OR "Lyncornis" OR "Nyctidromus" OR "Nyctiphrynus" OR "Nyctipolus" OR "Nyctiprogne" OR "Phalaenoptilus" OR "Setopagis" OR "Systellura" OR "Uropsalis" OR "Nyctibius" OR "Batrachostomus" OR "Podargus" OR "Steatornis" OR "Cariama" OR "Casuarius" OR "Dromaius" OR "Aethia" OR "Alca" OR "Alle" OR "Brachyramphus" OR "Cepphus" OR "Cerorhinca" OR "Fratercula" OR "Pinguinus" OR "Ptychoramphus" OR "Synthliboramphus" OR "Uria" OR "Burhinus" OR "Anarhynchus" OR "Charadrius" OR "Elseyornis" OR "Erythrogonys" OR "Hoploxypterus" OR "Oreopholus" OR "Peltohyas" OR "Phegornis" OR "Pluvialis" OR "Thinornis" OR "Vanellus" OR "Chionis" OR "Dromas" OR "Cursorius" OR "Glareola" OR "Rhinoptilus" OR "Stiltia" OR "Haematopus" OR "Actophilornis" OR "Hydrophasianus" OR

"Irediparra" OR "Jacana" OR "Metopidius" OR "Microparra" OR "Anous" OR "Chlidonias" OR "Chroicocephalus" OR "Creagrus" OR "Gelochelidon" OR "Gygis" OR "Hydrocoloeus" OR "Hydroprogne" OR "Ichthyaetus" OR "Larosterna" OR "Larus" OR "Leucophaeus" OR "Onychoprion" OR "Pagophila" OR "Phaetusa" OR "Rhodostethia" OR "Rissa" OR "Rynchops" OR "Sterna" OR "Sternula" OR "Thalasseus" OR "Xema" OR "Pedionomus" OR "Pluvianellus" OR "Pluvianus" OR "Himantopus" OR "Recurvirostra" OR "Nycticryphes" OR "Rostratula" OR "Actitis" OR "Arenaria" OR "Bartramia" OR "Calidris" OR "Coenocorypha" OR "Gallinago" OR "Limnodromus" OR "Limosa" OR "Lymnocryptes" OR "Numenius" OR "Phalaropus" OR "Scolopax" OR "Tringa" OR "Xenus" OR "Stercorarius" OR "Attagis" OR "Thinocorus" OR "Turnix" OR "Ciconia" OR "Ephippiorhynchus" OR "Jabiru" OR "Leptoptilos" OR "Mycteria" OR "Colius" OR "Urocolius" OR "Alectroenas" OR "Alopecoenas" OR "Caloenas" OR "Chalcophaps" OR "Claravis" OR "Columba" OR "Columbina" OR "Didunculus" OR "Drepanoptila" OR "Ducula" OR "Ectopistes" OR "Gallicolumba" OR "Geopelia" OR "Geophaps" OR "Geotrygon" OR "Goura" OR "Gymnophaps" OR "Hemiphaga" OR "Henicophaps" OR "Leptotila" OR "Leptotrygon" OR "Leucosarcia" OR "Lopholaimus" OR "Macropygia" OR "Metriopelia" OR "Ocyphaps" OR "Oena" OR "Otidiphaps" OR "Patagioenas" OR "Petrophassa" OR "Pezophaps" OR "Phapitreron" OR "Phaps" OR "Ptilinopus" OR "Raphus" OR "Reinwardtoena" OR "Spilopelia" OR "Streptopelia" OR "Treron" OR "Trugon" OR "Turacoena" OR "Turtur" OR "Uropelia" OR "Zenaida" OR "Zentrygon" OR "Actenoides" OR "Alcedo" OR "Caridonax" OR "Ceryle" OR "Ceyx" OR "Chloroceryle" OR "Cittura" OR "Corythornis" OR "Dacelo" OR "Halcyon" OR "Ispidina" OR "Lacedo" OR "Megaceryle" OR "Melidora" OR "Syma" OR "Tanysiptera" OR "Todiramphus" OR "Atelornis" OR "Brachypteracias" OR "Geobiastes" OR "Coracias" OR "Eurystomus" OR "Merops" OR "Nyctyornis" OR "Baryphthengus" OR "Electron" OR "Eumomota" OR "Hylomanes" OR "Momotus" OR "Todus" OR "Cacomantis" OR "Carpococcyx" OR "Centropus" OR "Cercococcyx" OR "Chrysococcyx" OR "Clamator" OR "Coccycua" OR "Coccyzus" OR "Coua" OR "Crotophaga" OR "Cuculus" OR "Dasylophus" OR "Dromococcyx" OR "Eudynamys" OR "Geococcyx" OR "Guira" OR "Hierococcyx" OR "Morococcyx" OR "Neomorphus" OR "Pachycoccyx" OR "Phaenicophaeus" OR "Piaya" OR "Rhinortha" OR "Scythrops" OR "Surniculus" OR "Tapera" OR "Urodynamis" OR

"Zanclostomus" OR "Dinornis" OR "Anomalopteryx" OR "Emeus" OR "Euryapteryx" OR "Eurypyga" OR "Rhynochetos" OR "Caracara" OR "Daptrius" OR "Falco" OR "Herpetotheres" OR "Ibycter" OR "Micrastur" OR "Microhierax" OR "Milvago" OR "Phalcoboenus" OR "Polihierax" OR "Spiziapteryx" OR "Aburria" OR "Chamaepetes" OR "Crax" OR "Mitu" OR "Nothocrax" OR "Oreophasis" OR "Ortalis" OR "Pauxi" OR "Penelope" OR "Penelopina" OR "Pipile" OR "Alectura" OR "Megapodius" OR "Acryllium" OR "Guttera" OR "Numida" OR "Callipepla" OR "Colinus" OR "Cyrtonyx" OR "Dendrortyx" OR "Odontophorus" OR "Oreortyx" OR "Philortyx" OR "Ptilopachus" OR "Rhynchortyx" OR "Alectoris" OR "Ammoperdix" OR "Arborophila" OR "Argusianus" OR "Bambusicola" OR "Bonasa" OR "Caloperdix" OR "Centrocercus" OR "Chrysolophus" OR "Coturnix" OR "Crossoptilon" OR "Dendragapus" OR "Excalfactoria" OR "Falcipennis" OR "Francolinus" OR "Gallus" OR "Haematortyx" OR "Ithaginis" OR "Lagopus" OR "Lerwa" OR "Lophophorus" OR "Lophura" OR "Lyrurus" OR "Meleagris" OR "Pavo" OR "Peliperdix" OR "Perdix" OR "Phasianus" OR "Polyplectron" OR "Pternistis" OR "Pucrasia" OR "Rhizothera" OR "Rollulus" OR "Scleroptila" OR "Syrmaticus" OR "Tetrao" OR "Tetraogallus" OR "Tetraophasis" OR "Tragopan" OR "Tympanuchus" OR "Gavia" OR "Aramus" OR "Anthropoides" OR "Antigone" OR "Balearica" OR "Bugeranus" OR "Grus" OR "Leucogeranus" OR "Heliornis" OR "Psophia" OR "Aenigmatolimnas" OR "Amaurolimnas" OR "Amaurornis" OR "Anurolimnas" OR "Aramides" OR "Atlantisia" OR "Coturnicops" OR "Crex" OR "Dryolimnas" OR "Eulabeornis" OR "Fulica" OR "Gallicrex" OR "Gallinula" OR "Gallirallus" OR "Laterallus" OR "Lewinia" OR "Micropygia" OR "Neocrex" OR "Nesoclopeus" OR "Paragallinula" OR "Pardirallus" OR "Porphyrio" OR "Porphyriops" OR "Porzana" OR "Rallicula" OR "Rallina" OR "Rallus" OR "Tribonyx" OR "Canirallus" OR "Sarothrura" OR "Leptosomus" OR "Corythaeola" OR "Corythaixoides" OR "Crinifer" OR "Musophaga" OR "Tauraco" OR "Opisthocomus" OR "Afrotis" OR "Ardeotis" OR "Chlamydotis" OR "Neotis" OR "Otis" OR "Tetrax" OR "Acanthisitta" OR "Pachyplichas" OR "Traversia" OR "Xenicus" OR "Acanthiza" OR "Acanthornis" OR "Aethomyias" OR "Aphelocephala" OR "Calamanthus" OR "Gerygone" OR "Hylacola" OR "Neosericornis" OR "Oreoscopus" OR "Origma" OR "Pachycare" OR "Pycnoptilus" OR "Pyrrholaemus" OR "Sericornis" OR "Smicrornis" OR "Acrocephalus" OR "Arundinax" OR "Calamonastides" OR "Hippolais" OR "Iduna" OR

"Nesillas" OR "Aegithalos" OR "Leptopoecile" OR "Psaltriparus" OR "Aegithina" OR "Alaemon" OR "Alauda" OR "Alaudala" OR "Ammomanes" OR "Calandrella" OR "Calendulauda" OR "Chersomanes" OR "Eremophila" OR "Eremopterix" OR "Galerida" OR "Lullula" OR "Melanocorypha" OR "Mirafra" OR "Pinarocorys" OR "Spizocorys" OR "Artamus" OR "Gymnorhina" OR "Melloria" OR "Bombycilla" OR "Buphagus" OR "Calcarius" OR "Plectrophenax" OR "Rhynchophanes" OR "Callaeas" OR "Heteralocha" OR "Philesturnus" OR "Calyptophilus" OR "Campephaga" OR "Ceblepyris" OR "Coracina" OR "Edolisoma" OR "Lalage" OR "Malindangia" OR "Pericrocotus" OR "Amaurospiza" OR "Cardinalis" OR "Caryothraustes" OR "Chlorothraupis" OR "Cyanocompsa" OR "Cyanoloxia" OR "Granatellus" OR "Habia" OR "Passerina" OR "Periporphyrus" OR "Pheucticus" OR "Piranga" OR "Spiza" OR "Certhia" OR "Salpornis" OR "Abroscopus" OR "Cettia" OR "Horornis" OR "Phyllergates" OR "Tesia" OR "Urosphena" OR "Chloropsis" OR "Cinclus" OR "Apalis" OR "Artisornis" OR "Camaroptera" OR "Cisticola" OR "Eremomela" OR "Heliolais" OR "Hypergerus" OR "Neomixis" OR "Oreolais" OR "Orthotomus" OR "Phragmacia" OR "Poliolais" OR "Prinia" OR "Schistolais" OR "Urolais" OR "Urorhipis" OR "Climacteris" OR "Cormobates" OR "Conopophaga" OR "Struthidea" OR "Aphelocoma" OR "Calocitta" OR "Cissa" OR "Coloeus" OR "Corvus" OR "Cyanocitta" OR "Cyanocorax" OR "Cyanolyca" OR "Cyanopica" OR "Dendrocitta" OR "Garrulus" OR "Gymnorhinus" OR "Nucifraga" OR "Perisoreus" OR "Pica" OR "Platylophus" OR "Podoces" OR "Psilorhinus" OR "Ptilostomus" OR "Pyrrhocorax" OR "Urocissa" OR "Ampelioides" OR "Ampelion" OR "Carpornis" OR "Cephalopterus" OR "Conioptilon" OR "Cotinga" OR "Doliornis" OR "Gymnoderus" OR "Haematoderus" OR "Lipaugus" OR "Perissocephalus" OR "Phoenicircus" OR "Phytotoma" OR "Pipreola" OR "Porphyrolaema" OR "Procnias" OR "Pyroderus" OR "Querula" OR "Rupicola" OR "Snowornis" OR "Xipholena" OR "Zaratornis" OR "Dasyornis" OR "Dicaeum" OR "Prionochilus" OR "Dicrurus" OR "Donacobius" OR "Dulus" OR "Emberiza" OR "Amadina" OR "Amandava" OR "Cryptospiza" OR "Erythrura" OR "Estrilda" OR "Euodice" OR "Euschistospiza" OR "Lagonosticta" OR "Lonchura" OR "Mandingoa" OR "Neochmia" OR "Nesocharis" OR "Nigrita" OR "Ortygospiza" OR "Parmoptila" OR "Pyrenestes" OR "Pytilia" OR "Spermophaga" OR "Stagonopleura" OR "Taeniopygia" OR "Uraeginthus" OR "Calyptomena" OR "Cymbirhynchus" OR "Eurylaimus" OR

"Psarisomus" OR "Serilophus" OR "Smithornis" OR "Chamaeza" OR "Formicarius" OR "Acanthis" OR "Agraphospiza" OR "Akialoa" OR "Bucanetes" OR "Carduelis" OR "Carpodacus" OR "Chloris" OR "Chlorodrepanis" OR "Chlorophonia" OR "Chrysocorythus" OR "Coccothraustes" OR "Crithagra" OR "Drepanis" OR "Eophona" OR "Euphonia" OR "Fringilla" OR "Haemorhous" OR "Hemignathus" OR "Himatione" OR "Leucosticte" OR "Linaria" OR "Linurgus" OR "Loxia" OR "Loxioides" OR "Loxops" OR "Magumma" OR "Melamprosops" OR "Oreomystis" OR "Paroreomyza" OR "Pinicola" OR "Procarduelis" OR "Pseudonestor" OR "Psittirostra" OR "Pyrrhoplectes" OR "Pyrrhula" OR "Rhodopechys" OR "Rhodospiza" OR "Serinus" OR "Spinus" OR "Telespiza" OR "Anabacerthia" OR "Anabazenops" OR "Ancistrops" OR "Anumbius" OR "Aphrastura" OR "Asthenes" OR "Automolus" OR "Berlepschia" OR "Campylorhamphus" OR "Certhiasomus" OR "Certhiaxis" OR "Cinclodes" OR "Clibanornis" OR "Coryphistera" OR "Cranioleuca" OR "Deconychura" OR "Dendrexetastes" OR "Dendrocincla" OR "Dendrocolaptes" OR "Dendroplex" OR "Drymornis" OR "Drymotoxeres" OR "Furnarius" OR "Geocerthia" OR "Geositta" OR "Glyphorynchus" OR "Heliobletus" OR "Hellmayrea" OR "Hylexetastes" OR "Lepidocolaptes" OR "Leptasthenura" OR "Limnoctites" OR "Limnornis" OR "Lochmias" OR "Margarornis" OR "Mazaria" OR "Megaxenops" OR "Metopothrix" OR "Microxenops" OR "Nasica" OR "Ochetorhynchus" OR "Phacellodomus" OR "Philydor" OR "Phleocryptes" OR "Premnoplex" OR "Premnornis" OR "Pseudasthenes" OR "Pseudocolaptes" OR "Pseudoseisura" OR "Pygarrhichas" OR "Roraimia" OR "Schoeniophylax" OR "Sclerurus" OR "Sittasomus" OR "Spartonoica" OR "Sylviorthorhynchus" OR "Synallaxis" OR "Syndactyla" OR "Tarphonomus" OR "Thripadectes" OR "Thripophaga" OR "Upucerthia" OR "Xenerpestes" OR "Xenops" OR "Xiphocolaptes" OR "Xiphorhynchus" OR "Grallaria" OR "Grallaricula" OR "Hylopezus" OR "Myrmothera" OR "Alopochelidon" OR "Atticora" OR "Cecropis" OR "Delichon" OR "Haplochelidon" OR "Hirundo" OR "Neochelidon" OR "Notiochelidon" OR "Orochelidon" OR "Petrochelidon" OR "Phedina" OR "Progne" OR "Psalidoprocne" OR "Pseudhirundo" OR "Pseudochelidon" OR "Ptyonoprogne" OR "Riparia" OR "Stelgidopteryx" OR "Tachycineta" OR "Hyliota" OR "Hypocolius" OR "Agelaioides" OR "Agelaius" OR "Agelasticus" OR "Amblycercus" OR "Amblyramphus" OR "Anumara" OR "Cacicus" OR "Chrysomus" OR "Curaeus" OR "Dives" OR "Dolichonyx" OR

"Euphagus" OR "Gnorimopsar" OR "Gymnomystax" OR "Hypopyrrhus" OR "Icterus" OR "Lampropsar" OR "Leistes" OR "Macroagelaius" OR "Molothrus" OR "Nesopsar" OR "Oreopsar" OR "Psarocolius" OR "Pseudoleistes" OR "Quiscalus" OR "Sturnella" OR "Xanthocephalus" OR "Xanthopsar" OR "Icteria" OR "Ifrita" OR "Irena" OR "Corvinella" OR "Eurocephalus" OR "Lanius" OR "Actinodura" OR "Argya" OR "Garrulax" OR "Grammatoptila" OR "Heterophasia" OR "Ianthocincla" OR "Laniellus" OR "Leiothrix" OR "Liocichla" OR "Minla" OR "Pterorhinus" OR "Trochalopteron" OR "Turdoides" OR "Bradypterus" OR "Cincloramphus" OR "Helopsaltes" OR "Locustella" OR "Megalurus" OR "Poodytes" OR "Machaerirhynchus" OR "Graueria" OR "Hylia" OR "Macrosphenus" OR "Melocichla" OR "Sylvietta" OR "Chlorophoneus" OR "Dryoscopus" OR "Laniarius" OR "Malaconotus" OR "Nilaus" OR "Rhodophoneus" OR "Tchagra" OR "Telophorus" OR "Amytornis" OR "Chenorhamphus" OR "Clytomyias" OR "Malurus" OR "Stipiturus" OR "Melampitta" OR "Melanocharis" OR "Oedistoma" OR "Toxorhamphus" OR "Melanopareia" OR "Acanthagenys" OR "Acanthorhynchus" OR "Anthochaera" OR "Anthornis" OR "Caligavis" OR "Entomyzon" OR "Epthianura" OR "Glycifohia" OR "Gymnomyza" OR "Lichmera" OR "Manorina" OR "Melidectes" OR "Melilestes" OR "Meliphaga" OR "Melipotes" OR "Melithreptus" OR "Myzomela" OR "Nesoptilotis" OR "Philemon" OR "Phylidonyris" OR "Prosthemadera" OR "Ptiloprora" OR "Ptilotula" OR "Pycnopygius" OR "Timeliopsis" OR "Xanthotis" OR "Menura" OR "Allenia" OR "Cinclocerthia" OR "Dumetella" OR "Margarops" OR "Melanoptila" OR "Melanotis" OR "Mimus" OR "Oreoscoptes" OR "Ramphocinclus" OR "Toxostoma" OR "Lamprospiza" OR "Mitrospingus" OR "Moho" OR "Mohoua" OR "Arses" OR "Carterornis" OR "Clytorhynchus" OR "Grallina" OR "Hypothymis" OR "Monarcha" OR "Myiagra" OR "Symposiachrus" OR "Terpsiphone" OR "Trochocercus" OR "Anthus" OR "Macronyx" OR "Motacilla" OR "Alethe" OR "Anthipes" OR "Brachypteryx" OR "Calliope" OR "Campicoloides" OR "Cercotrichas" OR "Chamaetylas" OR "Copsychus" OR "Cossypha" OR "Cyanoptila" OR "Cyornis" OR "Enicurus" OR "Erithacus" OR "Eumyias" OR "Ficedula" OR "Fraseria" OR "Irania" OR "Larvivora" OR "Leonardina" OR "Luscinia" OR "Melaenornis" OR "Monticola" OR "Muscicapa" OR "Myiomela" OR "Myioparus" OR "Myophonus" OR "Myrmecocichla" OR "Niltava" OR "Oenanthe" OR "Phoenicurus" OR "Pogonocichla" OR "Saxicola" OR "Sheppardia" OR "Sholicola" OR "Stiphrornis" OR "Tarsiger" OR

"Thamnolaea" OR "Vauriella" OR "Aethopyga" OR "Anabathmis" OR "Anthreptes" OR "Arachnothera" OR "Chalcomitra" OR "Cinnyris" OR "Cyanomitra" OR "Deleornis" OR "Hedydipna" OR "Kurochkinegramma" OR "Leptocoma" OR "Nectarinia" OR "Daphoenositta" OR "Nesospingus" OR "Nicator" OR "Notiomystis" OR "Aleadryas" OR "Oriolus" OR "Pitohui" OR "Sphecotheres" OR "Turnagra" OR "Colluricincla" OR "Falcunculus" OR "Melanorectes" OR "Pachycephala" OR "Pseudorectes" OR "Panurus" OR "Cicinnurus" OR "Diphyllodes" OR "Manucodia" OR "Paradisaea" OR "Ptiloris" OR "Paramythia" OR "Pardalotus" OR "Baeolophus" OR "Cephalopyrus" OR "Cyanistes" OR "Lophophanes" OR "Machlolophus" OR "Melaniparus" OR "Melanochlora" OR "Pardaliparus" OR "Parus" OR "Periparus" OR "Poecile" OR "Pseudopodoces" OR "Sittiparus" OR "Sylviparus" OR "Basileuterus" OR "Cardellina" OR "Catharopeza" OR "Dendroica" OR "Geothlypis" OR "Helmitheros" OR "Leiothlypis" OR "Limnothlypis" OR "Mniotilta" OR "Myioborus" OR "Myiothlypis" OR "Oporornis" OR "Oreothlypis" OR "Parkesia" OR "Protonotaria" OR "Seiurus" OR "Setophaga" OR "Vermivora" OR "Aimophila" OR "Ammodramus" OR "Amphispiza" OR "Arremon" OR "Arremonops" OR "Artemisiospiza" OR "Atlapetes" OR "Calamospiza" OR "Chlorospingus" OR "Chondestes" OR "Junco" OR "Melospiza" OR "Melozone" OR "Oreothraupis" OR "Oriturus" OR "Passerculus" OR "Passerella" OR "Peucaea" OR "Pezopetes" OR "Pipilo" OR "Pooecetes" OR "Pselliophorus" OR "Rhynchospiza" OR "Spizella" OR "Spizelloides" OR "Xenospiza" OR "Zonotrichia" OR "Carpospiza" OR "Gymnoris" OR "Hypocryptadius" OR "Montifringilla" OR "Onychostruthus" OR "Passer" OR "Petronia" OR "Pyrgilauda" OR "Alcippe" OR "Gampsorhynchus" OR "Illadopsis" OR "Jabouilleia" OR "Kenopia" OR "Laticilla" OR "Malacocincla" OR "Malacopteron" OR "Napothera" OR "Pellorneum" OR "Rimator" OR "Trichastoma" OR "Amalocichla" OR "Drymodes" OR "Eopsaltria" OR "Eugerygone" OR "Heteromyias" OR "Melanodryas" OR "Microeca" OR "Monachella" OR "Pachycephalopsis" OR "Peneoenanthe" OR "Peneothello" OR "Petroica" OR "Poecilodryas" OR "Tregellasia" OR "Peucedramus" OR "Microligea" OR "Phaenicophilus" OR "Xenoligea" OR "Phylloscopus" OR "Picathartes" OR "Antilophia" OR "Ceratopipra" OR "Chiroxiphia" OR "Chloropipo" OR "Corapipo" OR "Cryptopipo" OR "Heterocercus" OR "Ilicura" OR "Lepidothrix" OR "Machaeropterus" OR "Manacus" OR "Masius" OR "Neopelma" OR "Pipra" OR "Pseudopipra" OR "Tyranneutes" OR

"Xenopipo" OR "Erythropitta" OR "Hydrornis" OR "Pitta" OR "Batis" OR "Platysteira" OR "Anaplectes" OR "Bubalornis" OR "Euplectes" OR "Malimbus" OR "Plocepasser" OR "Ploceus" OR "Quelea" OR "Sporopipes" OR "Pnoepyga" OR "Microbates" OR "Polioptila" OR "Ramphocaenus" OR "Garritornis" OR "Prunella" OR "Cinclosoma" OR "Ptilorrhoa" OR "Phainopepla" OR "Phainoptila" OR "Ptiliogonys" OR "Ailuroedus" OR "Amblyornis" OR "Ptilonorhynchus" OR "Sericulus" OR "Alophoixus" OR "Andropadus" OR "Arizelocichla" OR "Atimastillas" OR "Baeopogon" OR "Bleda" OR "Chlorocichla" OR "Criniger" OR "Eurillas" OR "Hemixos" OR "Hypsipetes" OR "Iole" OR "Ixos" OR "Neolestes" OR "Phyllastrephus" OR "Pycnonotus" OR "Spizixos" OR "Stelgidillas" OR "Tricholestes" OR "Regulus" OR "Anthoscopus" OR "Auriparus" OR "Remiz" OR "Rhagologus" OR "Eleoscytalopus" OR "Liosceles" OR "Psilorhamphus" OR "Pteroptochos" OR "Scelorchilus" OR "Scytalopus" OR "Teledromas" OR "Rhipidura" OR "Rhodinocichla" OR "Sapayoa" OR "Sitta" OR "Spindalis" OR "Chelidorhynx" OR "Culicicapa" OR "Elminia" OR "Acridotheres" OR "Agropsar" OR "Ampeliceps" OR "Aplonis" OR "Basilornis" OR "Cinnyricinclus" OR "Creatophora" OR "Gracula" OR "Gracupica" OR "Hartlaubius" OR "Hylopsar" OR "Lamprotornis" OR "Leucopsar" OR "Mino" OR "Notopholia" OR "Onychognathus" OR "Pastor" OR "Poeoptera" OR "Rhabdornis" OR "Sarcops" OR "Scissirostrum" OR "Speculipastor" OR "Spodiopsar" OR "Streptocitta" OR "Sturnia" OR "Sturnus" OR "Chamaea" OR "Chleuasicus" OR "Chloropeta" OR "Chrysomma" OR "Conostoma" OR "Fulvetta" OR "Lioparus" OR "Paradoxornis" OR "Pseudoalcippe" OR "Psittiparus" OR "Rhopophilus" OR "Sinosuthora" OR "Suthora" OR "Sylvia" OR "Teretistris" OR "Akletos" OR "Ampelornis" OR "Aprositornis" OR "Batara" OR "Cercomacra" OR "Cercomacroides" OR "Cymbilaimus" OR "Dichrozona" OR "Drymophila" OR "Dysithamnus" OR "Epinecrophylla" OR "Euchrepomis" OR "Formicivora" OR "Frederickena" OR "Gymnocichla" OR "Gymnopithys" OR "Hafferia" OR "Herpsilochmus" OR "Hylophylax" OR "Hypocnemis" OR "Hypocnemoides" OR "Hypoedaleus" OR "Isleria" OR "Mackenziaena" OR "Megastictus" OR "Microrhopias" OR "Myrmeciza" OR "Myrmelastes" OR "Myrmoborus" OR "Myrmochanes" OR "Myrmoderus" OR "Myrmophylax" OR "Myrmorchilus" OR "Myrmornis" OR "Myrmotherula" OR "Neoctantes" OR "Oneillornis" OR "Percnostola" OR "Phaenostictus" OR "Phlegopsis" OR "Pithys" OR "Poliocrania" OR "Pygiptila" OR "Pyriglena" OR "Rhegmatorhina" OR

"Rhopias" OR "Rhopornis" OR "Sakesphorus" OR "Sciaphylax" OR "Sclateria" OR "Sipia" OR "Taraba" OR "Thamnistes" OR "Thamnomanes" OR "Thamnophilus" OR "Willisornis" OR "Xenornis" OR "Anisognathus" OR "Bangsia" OR "Buthraupis" OR "Calochaetes" OR "Catamblyrhynchus" OR "Catamenia" OR "Charitospiza" OR "Chlorochrysa" OR "Chlorophanes" OR "Chlorornis" OR "Chrysothlypis" OR "Cissopis" OR "Cnemoscopus" OR "Coereba" OR "Compsospiza" OR "Compsothraupis" OR "Conirostrum" OR "Conothraupis" OR "Coryphospingus" OR "Creurgops" OR "Cyanerpes" OR "Cyanicterus" OR "Cypsnagra" OR "Dacnis" OR "Delothraupis" OR "Diglossa" OR "Diuca" OR "Dolospingus" OR "Donacospiza" OR "Dubusia" OR "Emberizoides" OR "Embernagra" OR "Eucometis" OR "Euneornis" OR "Geospiza" OR "Gubernatrix" OR "Haplospiza" OR "Hemispingus" OR "Hemithraupis" OR "Heterospingus" OR "Incaspiza" OR "Iridophanes" OR "Iridosornis" OR "Lanio" OR "Lophospingus" OR "Loxigilla" OR "Loxipasser" OR "Melanodera" OR "Melanospiza" OR "Melopyrrha" OR "Nemosia" OR "Nephelornis" OR "Oreomanes" OR "Oryzoborus" OR "Parkerthraustes" OR "Paroaria" OR "Phrygilus" OR "Piezorina" OR "Pipraeidea" OR "Poospiza" OR "Pyrrhocoma" OR "Ramphocelus" OR "Rhodospingus" OR "Saltator" OR "Saltatricula" OR "Schistochlamys" OR "Sericossypha" OR "Sicalis" OR "Sporophila" OR "Stephanophorus" OR "Tachyphonus" OR "Tangara" OR "Tersina" OR "Thlypopsis" OR "Thraupis" OR "Tiaris" OR "Trichothraupis" OR "Volatinia" OR "Wetmorethraupis" OR "Xenodacnis" OR "Xenospingus" OR "Tichodroma" OR "Macronus" OR "Pomatorhinus" OR "Spelaeornis" OR "Stachyridopsis" OR "Stachyris" OR "Iodopleura" OR "Laniisoma" OR "Laniocera" OR "Myiobius" OR "Onychorhynchus" OR "Oxyruncus" OR "Pachyramphus" OR "Schiffornis" OR "Terenotriccus" OR "Tityra" OR "Xenopsaris" OR "Campylorhynchus" OR "Cantorchilus" OR "Catherpes" OR "Cinnycerthia" OR "Cistothorus" OR "Cyphorhinus" OR "Henicorhina" OR "Microcerculus" OR "Odontorchilus" OR "Pheugopedius" OR "Salpinctes" OR "Thryomanes" OR "Thryophilus" OR "Thryothorus" OR "Troglodytes" OR "Uropsila" OR "Catharus" OR "Cichlopsis" OR "Entomodestes" OR "Geokichla" OR "Hylocichla" OR "Ixoreus" OR "Myadestes" OR "Neocossyphus" OR "Ridgwayia" OR "Sialia" OR "Stizorhina" OR "Turdus" OR "Zoothera" OR "Agriornis" OR "Alectrurus" OR "Anairetes" OR "Aphanotriccus" OR "Arundinicola" OR "Atalotriccus" OR "Attila" OR "Camptostoma" OR "Capsiempis" OR

"Casiornis" OR "Cnemarchus" OR "Cnemotriccus" OR "Cnipodectes" OR "Colonia" OR "Colorhamphus" OR "Conopias" OR "Contopus" OR "Corythopis" OR "Culicivora" OR "Deltarhynchus" OR "Elaenia" OR "Empidonax" OR "Empidonomus" OR "Euscarthmus" OR "Fluvicola" OR "Griseotyrannus" OR "Gubernetes" OR "Hemitriccus" OR "Heteroxolmis" OR "Hirundinea" OR "Hymenops" OR "Inezia" OR "Knipolegus" OR "Lathrotriccus" OR "Legatus" OR "Leptopogon" OR "Lessonia" OR "Lophotriccus" OR "Machetornis" OR "Mecocerculus" OR "Megarynchus" OR "Mionectes" OR "Mitrephanes" OR "Muscigralla" OR "Muscisaxicola" OR "Myiarchus" OR "Myiodynastes" OR "Myiopagis" OR "Myiophobus" OR "Myiornis" OR "Myiotheretes" OR "Myiotriccus" OR "Myiozetetes" OR "Neopipo" OR "Neoxolmis" OR "Nephelomyias" OR "Ochthoeca" OR "Ochthornis" OR "Oncostoma" OR "Ornithion" OR "Phaeomyias" OR "Philohydor" OR "Phyllomyias" OR "Phylloscartes" OR "Piprites" OR "Pitangus" OR "Platyrinchus" OR "Poecilotriccus" OR "Pogonotriccus" OR "Polioxolmis" OR "Polystictus" OR "Pseudelaenia" OR "Pseudocolopteryx" OR "Pseudotriccus" OR "Pyrocephalus" OR "Pyrrhomyias" OR "Ramphotrigon" OR "Rhynchocyclus" OR "Rhytipterna" OR "Satrapa" OR "Sayornis" OR "Serpophaga" OR "Silvicultrix" OR "Sirystes" OR "Stigmatura" OR "Sublegatus" OR "Suiriri" OR "Tachuris" OR "Taeniotriccus" OR "Todirostrum" OR "Tolmomyias" OR "Tumbezia" OR "Tyrannopsis" OR "Tyrannulus" OR "Tyrannus" OR "Uromyias" OR "Xenotriccus" OR "Xolmis" OR "Zimmerius" OR "Urocynchramus" OR "Artamella" OR "Calicalicus" OR "Cyanolanius" OR "Euryceros" OR "Falculea" OR "Hypositta" OR "Leptopterus" OR "Mystacornis" OR "Newtonia" OR "Oriolia" OR "Philentoma" OR "Prionops" OR "Pseudobias" OR "Schetba" OR "Tephrodornis" OR "Tylas" OR "Vanga" OR "Xenopirostris" OR "Anomalospiza" OR "Vidua" OR "Cyclarhis" OR "Erpornis" OR "Hylophilus" OR "Pteruthius" OR "Vireo" OR "Vireolanius" OR "Zeledonia" OR "Apalopteron" OR "Dasycrotapha" OR "Heleia" OR "Lophozosterops" OR "Sterrhoptilus" OR "Yuhina" OR "Zosterops" OR "Zosterornis" OR "Agamia" OR "Ardea" OR "Ardeola" OR "Botaurus" OR "Bubulcus" OR "Butorides" OR "Cochlearius" OR "Dupetor" OR "Egretta" OR "Gorsachius" OR "Ixobrychus" OR "Nyctanassa" OR "Nycticorax" OR "Pilherodius" OR "Syrigma" OR "Tigriornis" OR "Tigrisoma" OR "Zebrilus" OR "Balaeniceps" OR "Pelecanus" OR "Scopus" OR "Bostrychia" OR "Eudocimus" OR "Geronticus" OR "Lophotibis" OR "Mesembrinibis" OR "Nipponia" OR "Phimosus"

OR "Platalea" OR "Plegadis" OR "Pseudibis" OR "Theristicus" OR "Threskiornis" OR "Phaethon" OR "Phoeniconaias" OR "Phoenicoparrus" OR "Phoenicopterus" OR "Bucco" OR "Chelidoptera" OR "Hapaloptila" OR "Hypnelus" OR "Malacoptila" OR "Micromonacha" OR "Monasa" OR "Nonnula" OR "Notharchus" OR "Nystalus" OR "Capito" OR "Eubucco" OR "Brachygalba" OR "Galbalcyrhynchus" OR "Galbula" OR "Jacamaralcyon" OR "Jacamerops" OR "Indicator" OR "Melichneutes" OR "Melignomon" OR "Prodotiscus" OR "Buccanodon" OR "Gymnobucco" OR "Lybius" OR "Pogoniulus" OR "Stactolaema" OR "Trachyphonus" OR "Psilopogon" OR "Blythipicus" OR "Campephilus" OR "Campethera" OR "Celeus" OR "Chloropicus" OR "Chrysocolaptes" OR "Chrysophlegma" OR "Colaptes" OR "Dendrocopos" OR "Dendrocoptes" OR "Dendropicos" OR "Dinopium" OR "Dryobates" OR "Dryocopus" OR "Gecinulus" OR "Geocolaptes" OR "Jynx" OR "Leiopicus" OR "Leuconotopicus" OR "Meiglyptes" OR "Melanerpes" OR "Micropternus" OR "Mulleripicus" OR "Nesoctites" OR "Picoides" OR "Piculus" OR "Picumnus" OR "Picus" OR "Reinwardtipicus" OR "Sasia" OR "Sphyrapicus" OR "Veniliornis" OR "Yungipicus" OR "Andigena" OR "Aulacorhynchus" OR "Pteroglossus" OR "Ramphastos" OR "Selenidera" OR "Semnornis" OR "Aechmophorus" OR "Podiceps" OR "Podilymbus" OR "Poliocephalus" OR "Rollandia" OR "Tachybaptus" OR "Diomedea" OR "Phoebastria" OR "Phoebetria" OR "Thalassarche" OR "Hydrobates" OR "Oceanodroma" OR "Fregetta" OR "Garrodia" OR "Nesofregetta" OR "Oceanites" OR "Pelagodroma" OR "Aphrodroma" OR "Ardenna" OR "Bulweria" OR "Calonectris" OR "Daption" OR "Fulmarus" OR "Halobaena" OR "Macronectes" OR "Pachyptila" OR "Pagodroma" OR "Pelecanoides" OR "Procellaria" OR "Pseudobulweria" OR "Pterodroma" OR "Puffinus" OR "Thalassoica" OR "Cacatua" OR "Callocephalon" OR "Calyptorhynchus" OR "Eolophus" OR "Lophochroa" OR "Nymphicus" OR "Probosciger" OR "Alipiopsitta" OR "Amazona" OR "Anodorhynchus" OR "Ara" OR "Aratinga" OR "Bolborhynchus" OR "Brotogeris" OR "Conuropsis" OR "Cyanoliseus" OR "Cyanopsitta" OR "Deroptyus" OR "Diopsittaca" OR "Enicognathus" OR "Eupsittula" OR "Forpus" OR "Graydidascalus" OR "Guarouba" OR "Guaruba" OR "Hapalopsittaca" OR "Leptosittaca" OR "Myiopsitta" OR "Nannopsittaca" OR "Orthopsittaca" OR "Pionites" OR "Pionopsitta" OR "Pionus" OR "Poicephalus" OR "Primolius" OR "Psilopsiagon" OR "Psittacara" OR "Psittacus" OR "Pyrilia" OR "Pyrrhura" OR

"Rhynchopsitta" OR "Thectocercus" OR "Touit" OR "Triclaria" OR "Agapornis" OR "Alisterus" OR "Aprosmictus" OR "Barnardius" OR "Bolbopsittacus" OR "Chalcopsitta" OR "Charmosyna" OR "Coracopsis" OR "Cyanoramphus" OR "Cyclopsitta" OR "Eclectus" OR "Eos" OR "Eunymphicus" OR "Geoffroyus" OR "Glossopsitta" OR "Lathamus" OR "Loriculus" OR "Lorius" OR "Melopsittacus" OR "Micropsitta" OR "Neophema" OR "Neopsephotus" OR "Neopsittacus" OR "Northiella" OR "Oreopsittacus" OR "Parvipsitta" OR "Pezoporus" OR "Phigys" OR "Platycercus" OR "Polytelis" OR "Prioniturus" OR "Prosopeia" OR "Psephotellus" OR "Psephotus" OR "Pseudeos" OR "Psittacella" OR "Psittacula" OR "Psittaculirostris" OR "Psitteuteles" OR "Psittinus" OR "Psittrichas" OR "Purpureicephalus" OR "Tanygnathus" OR "Trichoglossus" OR "Vini" OR "Nestor" OR "Strigops" OR "Pterocles" OR "Syrrhaptes" OR "Rhea" OR "Aptenodytes" OR "Eudyptes" OR "Eudyptula" OR "Megadyptes" OR "Pygoscelis" OR "Spheniscus" OR "Aegolius" OR "Asio" OR "Athene" OR "Bubo" OR "Glaucidium" OR "Ketupa" OR "Lophostrix" OR "Megascops" OR "Micrathene" OR "Ninox" OR "Otus" OR "Pseudoscops" OR "Psiloscops" OR "Pulsatrix" OR "Sceloglaux" OR "Scotopelia" OR "Strix" OR "Surnia" OR "Xenoglaux" OR "Phodilus" OR "Tyto" OR "Aepyornis" OR "Mullerornis" OR "Struthio" OR "Anhinga" OR "Fregata" OR "Leucocarbo" OR "Microcarbo" OR "Nannopterum" OR "Phalacrocorax" OR "Morus" OR "Sula" OR "Crypturellus" OR "Eudromia" OR "Nothocercus" OR "Nothoprocta" OR "Nothura" OR "Rhynchotus" OR "Tinamotis" OR "Tinamus" OR "Apalharpactes" OR "Apaloderma" OR "Harpactes" OR "Pharomachrus" OR "Trogon" OR "Tetrastes" OR "Hesperiphona") AND ("laying date" OR "lay date" OR "first egg" OR "clutch size" OR "eggs laid" OR "number of eggs" OR "fledgling*" OR "fledging" OR "reproductive success" OR "fitness"))

### References in Supplementary Materials:

Dhondt, A.A., Eyckerman, R., Moermans, R. & Hublé, J. (1984). Habitat and laying date of Great and Blue Tit Parus major and P. caeruleus. *Ibis*, 126, 388–397.

Nakagawa, S., Lagisz, M., Jennions, M.D., Koricheva, J., Noble, D.W.A., Parker, T.H., *et al.* (2022). Methods for testing publication bias in ecological and evolutionary meta-analyses. *Methods in Ecology and Evolution*, 13, 4–21.

Nakagawa, S., Lagisz, M., O'Dea, R.E., Rutkowska, J., Yang, Y., Noble, D.W.A., *et al.* (2021). The orchard plot: Cultivating a forest plot for use in ecology, evolution, and beyond. *Research Synthesis Methods*, 12, 4–12.
